# Multimodal MRI prediction of cognitive functioning across the lifespan: separating between-person differences from within-person changes

**DOI:** 10.64898/2026.03.15.711457

**Authors:** Kseniia Konopkina, Irina Buianova, Farzane Lal Khakpoor, Sunthud Pornprasertmanit, Micaela Chan, Narun Pat

## Abstract

Brain MRI shows promise for predicting cognitive functioning, but its utility depends on its capacity to capture stable between-person differences (e.g., patient stratification), longitudinal within-person changes (e.g., prognosis, treatment monitoring), or both. Using longitudinal data from 450 adults (aged 21–90; up to three waves, five years apart) in the Dallas Lifespan Brain Study, we benchmarked five modalities, task fMRI, functional connectivity (FC), structural MRI (sMRI), diffusion-weighted imaging (DWI), and arterial spin labeling (ASL), across 37 phenotypes and their combination. Stacking all MRI modalities into one marker predicted cognitive functioning with the highest accuracy (R²=.51), followed by DWI and FC. Variance decomposition showed MRI markers explained substantial between-person variance (up to 60.3%) but modest within-person changes (up to 17.2%) in cognitive functioning. Commonality analysis revealed most markers, except ASL, overlapped with age-related variance in cognitive functioning. These findings clarify the strengths and limitations of MRI markers for stratifying and monitoring cognitive aging.

---

For decades, cognitive neuroscientists have examined how age-related differences in cognitive functioning are associated with variations in brain structure and function, primarily using MRI data (see reviews ^1,2^). More recently, the integration of machine-learning frameworks has shifted the field from describing associations to generating out-of-sample predictions, ^3^ namely, predicting cognitive performance in previously unseen individuals using patterns learned from an independent training sample ^4^.

Evaluating model performance on individuals who were not involved in model development provides a direct test of the generalizability of MRI-based markers of cognitive functioning and, importantly, clarifies the extent to which different MRI modalities capture information relevant to cognitive variability ^5^.

Despite these advances, most predictive studies remain cross-sectional, leaving unclear the extent to which MRI markers capture stable between-person differences versus longitudinal within-person changes over time. Distinguishing these two sources of cognitive variation is crucial, as cognitive functioning varies not only between individuals in a relatively stable manner ^6^, but also changes within individuals over time ^7^. Understanding the capacity of the MRI markers is essential for setting realistic expectations about their potential applications. As an analogy, markers reflecting stable between-person differences may be valuable for diagnostic purposes, identifying individuals with consistently high or low levels of cognitive functioning. In contrast, markers sensitive to within-person changes would be more suitable for prognostic use, such as forecasting trajectories of cognitive decline. And because MRIs of different modalities reflect different aspects of the brain ^8^, the extent to which markers from various modalities can fulfil these complementary roles remains an open question.

Early machine-learning efforts focused on single-modality predictors, primarily using functional connectivity (FC) from resting-state functional MRI ^9–14^ or brain morphometry from structural MRI (sMRI) ^15–17^ to predict cognitive functioning. These unimodal approaches yielded mixed and generally modest prediction accuracies across different cognitive domains. Acknowledging that distinct neuroimaging modalities capture complementary aspects of the brain ^8^, recent studies have increasingly adopted multimodal approaches ^18–22^. One such strategy is stacking, in which machine learning models are trained separately for each modality, and their predicted values are subsequently combined as input features for a second-level model that generates the final prediction ^18,23^. Multimodal stacking usually improves the predictive accuracy of cognition beyond that achievable by any single modality, suggesting its ability to capture the complex interplay between various levels of brain organisation ^3,19,20,24,25^. These gains in predictive accuracy, however, do not in themselves clarify whether the models are capturing stable between-person differences or dynamic within-person changes throughout the aging process.

A cross-sectional design offers different perspectives on cognitive ageing than a longitudinal one. Cross-sectional studies, which compare people of different ages, have consistently reported an association between age and cognitive decline ^2,7,26–29^.

Longitudinal research has challenged the notion of inevitable decline by demonstrating marked heterogeneity in age-related cognitive change. Some individuals maintain stable performance well into late adulthood, while others show improvement or decline at varying rates ^28,30–35^. This fundamental discrepancy highlights the need to identify MRI markers that can account for both the general trends observed between people and the diverse patterns of change that occur within them over time.

The present study addresses this gap using longitudinal data from the Dallas Lifespan Brain Study (DLBS) ^36^ to benchmark the capacity of MRI markers of different modalities in capturing within-vs between-person variations in cognitive functioning. At baseline, participants were 21 to 89 years old and underwent repeated MRI scans at five-year intervals, up to three waves in total. We developed machine learning models to predict cognitive functioning using 37 neuroimaging phenotypes, spanning five MRI modalities: (1) task-based functional MRI (task-fMRI), which reflects BOLD activity associated with task events; (2) resting-state and task-based functional connectivity (FC), which captures correlations in BOLD time series across regions; (3) structural MRI (sMRI), which measures brain morphology; (4) diffusion-weighted imaging (DWI), which quantifies water diffusion patterns to infer structural connectivity and tractometry; and (5) arterial spin labeling (ASL), which estimates cerebral blood flow. Models were first trained on individual neuroimaging phenotypes (phenotype-specific models), then on combined phenotypes within each modality (modality-specific stacking models), and finally across all modalities (multimodal stacking models). These models aimed to predict out-of-sample cognitive functioning, operationalised as a general factor of cognition (g-score) derived from a higher-order g-factor model ^37^ using exploratory structural equation modelling (ESEM).

Our study pursued three primary objectives. First, we aimed to estimate and compare the accuracy of predicting cognitive functioning from individual neuroimaging phenotypes and their combinations. Second, we sought to quantify the extent to which MRI markers derived from various neuroimaging phenotypes accounted for within-vs. between-person variation in cognitive functioning. To achieve this, we employed linear mixed-effects models to separate within-person from between-person variance of cognitive functioning and assessed how much each MRI marker explained each type of variance of cognitive functioning via variance decomposition ^38^. Third, acknowledging that cognitive functioning changes with age, we aimed to determine how much of the age-related variance in cognitive functioning could be explained by these MRI markers; this was addressed using commonality analyses. Figure 1 illustrates our overall analytical framework.

**Figure 1.**
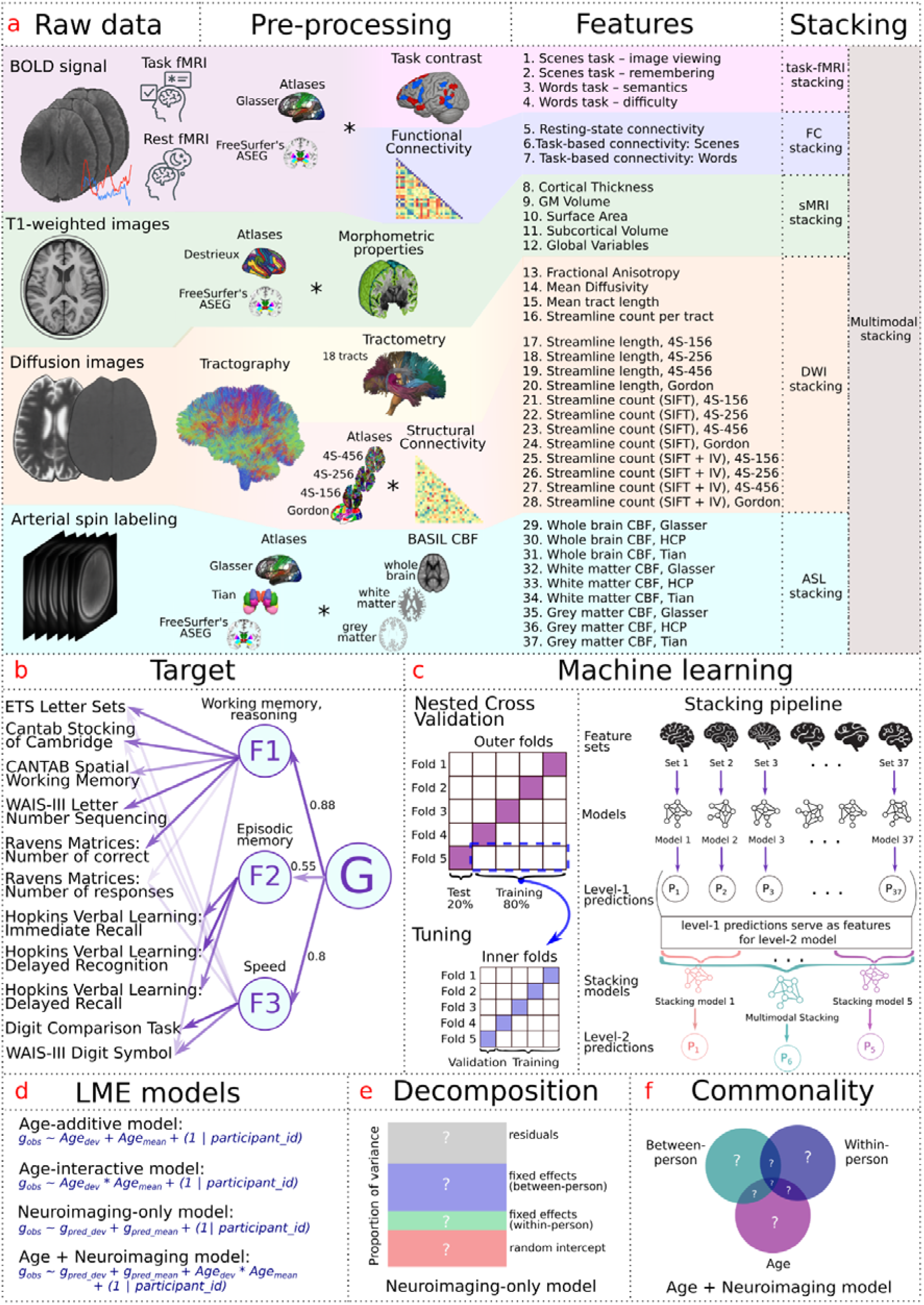
The schematic representation of the study. a. **Data to features.** Raw BOLD, T1-weighted, diffusion, and arterial spin labelling scans from the DLBS dataset were pre-processed to extract 37 neuroimaging feature sets, covering task-based functional MRI (task-fMRI), which reflects BOLD activity associated with task events; resting-state and task-based functional connectivity (FC), which captures correlations in BOLD time series across regions; structural MRI (sMRI), which measures brain morphology; diffusion-weighted imaging (DWI) which quantifies water diffusion patterns to infer structural connectivity and tractometry; and arterial spin labeling (ASL), which estimates cerebral blood flow. b. **Target.** The scheme displays the results of the Exploratory Structural Equation Modeling (ESEM). Cognitive functioning was estimated as a higher-order g-score that loads on three latent factors, underlying working-memory & reasoning, episodic memory, and processing speed, estimated from a battery of 11 cognitive measurements. G-score served as the prediction target. c. **Machine-learning workflow.** Each feature set entered a nested five-fold cross-validation. In the outer loop, participants were split into five folds with all waves from the same participant kept together to prevent leakage; one fold was held out for testing while the remainder formed the training set. The inner folds were used for hyperparameter tuning. We trained phenotype-specific models to predict the g-score using each neuroimaging feature set, resulting in level-1 predictions. These level-1 predictions served as inputs for six level-2 stacking models. The level-2 stacking models included five modality-specific stacking models, based on each of the five modalities, and one multimodal stacking model, based on the combination across all phenotypes. d. **Linear mixed-effects models.** Four linear mixed-effects models were applied to the outer-loop test-set predictions: age-additive, age-interactive, neuroimaging-only, and age-plus-neuroimaging. In all models, the observed g-score was the outcome. Age was decomposed into age*_mean_*, capturing between-person age differences, and age*_dev_*, capturing within-person age changes. MRI predictions were decomposed into g*_pred_mean_*, capturing between-person MRI markers, and g*_pred_dev_*, capturing within-person MRI markers. The neuroimaging-only model was used for variance decomposition, and the age-plus-neuroimaging model was used for commonality analysis. e. **Decomposition analysis.** Variance of cognitive functioning (g_obs_) as explained by the neuroimaging-only model is decomposed into within-person and between-person variance. We used this analysis to quantify how much the within-person MRI marker (g_pred_dev_) explained the within-person variance of cognitive functioning, and similarly, how much the between-person MRI marker (g_pred_mean_) explained the between-person variance of cognitive functioning. f. **Commonality analysis.** The analysis partitions explained variance into components shared by age and MRI predictions and components unique to each, evaluated at the within-person and between-person levels. This is to quantify the extent to which MRI markers accounted for age-related variation in cognitive functioning at within-and between-person levels. **Abbreviations:** DWI – diffusion-weighted imaging; sMRI – structural magnetic resonance imaging; task-fMRI – task-based functional magnetic resonance imaging; FC – functional connectivity; ASL – arterial spin labelling; CBF – cerebral blood flow; SIFT2 – Spherical-deconvolution Informed Filtering of Tractograms; 4S-***p (e.g., 4S-156p) – Schaefer atlas with *** parcels; HCP – Human Connectome Project parcellation that includes a combination of Glasser and ASEG atlases.

## Results

### Derivation of general cognitive ability (g-score)

Our primary measure of cognitive functioning was a g-score, derived using a hierarchical Exploratory Structural Equation Model (ESEM) ^39^. A parallel analysis indicated that three first-order factors adequately captured the latent structure of the cognitive battery. The factor-loading patterns reflected interpretable groupings: a working-memory/reasoning factor, an episodic-memory factor, and a processing-speed factor. Model fit was strong across folds, with RMSEA ranging from 0.046 to 0.056, CFI from 0.985 to 0.990, SRMR from 0.018 to 0.023, and TLI consistently exceeding 0.96. The intraclass correlation coefficient for g-score was 0.88, indicating that 88% of the variance occurred between individuals and 12% within individuals over time. Overall, the g-score accounted for approximately 58% to 62% of the variance in observed cognitive scores. Individual scores on the higher-order g factor were estimated from the fitted ESEM using the regression method. We extracted this factor score, referred to as the “observed g-score,” and used it as the target variable in our MRI-based machine-learning models. Full details appear in Supplementary Tables 1 and 2.

### The relationship between g-score and age

We examined the association between the observed g-score and age (Figure 2a). The age-additive model regressed the observed g-score on age*_mean_* (each individual’s average age across waves, capturing between-person age effects) and age*_dev_* (the deviation from age*_mean_* at each wave, capturing within-person age effects). The age-interactive model added the interaction between age*_mean_* and age*_dev_* to this baseline model, yielding a significant improvement in fit (χ²(1) = 59.65, p <.001) and explaining nearly half of the marginal R² (53%) in observed g-scores. As shown in Figure 2c, the interaction indicated that within-person age-related cognitive decline was steeper among older adults than younger adults.

**Figure 2.**
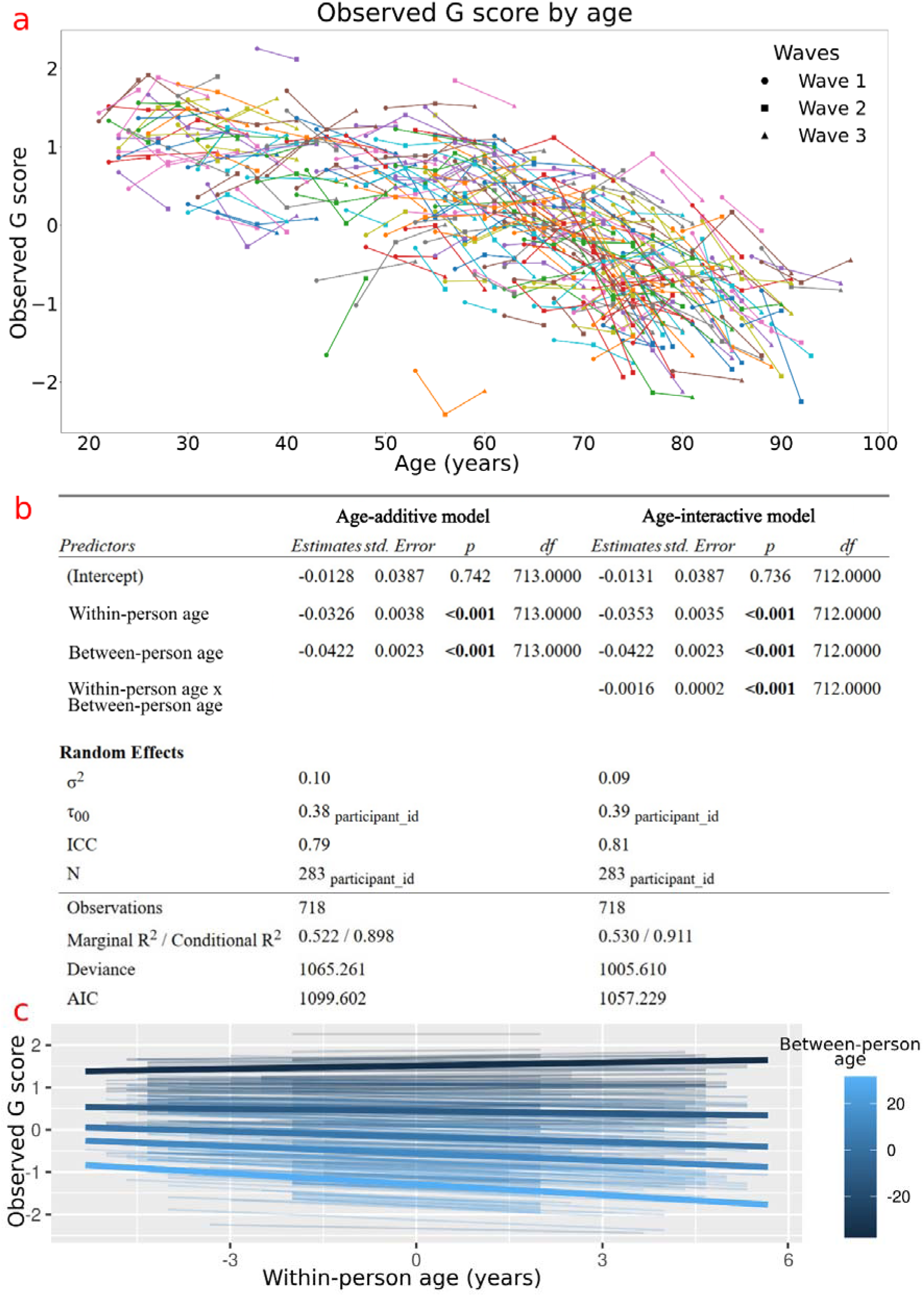
Relationship between age and observed g-score. **a. Observed cognitive g-score as a function of age across waves.** The observed g-score is a proxy for general cognitive functioning, derived from 11 cognitive measurements using hierarchical exploratory structural equation modelling (ESEM). Each point represents an assessment from one participant, with solid lines connecting repeated observations from the same individual over up to three waves collected roughly four to five years apart. Colours distinguish individuals only to facilitate visualisation, and marker shapes indicate the wave. Each line, therefore, traces a participant’s longitudinal trajectory. The dispersion of individual trajectories highlights substantial heterogeneity in cognitive ageing: while many participants decline, others remain stable, and some even improve. This observed g-score served as the target variable for all machine-learning models predicting g-score from multimodal MRI features. Note: the figure includes only participants with data from at least two waves to illustrate longitudinal trends. **b. Linear mixed-effects model results for the association between observed g-score and age.** The age-additive model included within-person (age*_dev_*) and between-person (age*_mean_*) age variables, whereas the age-interactive model additionally included the interaction between these age variables. Reported are fixed and random effects, model fit indices, and variance explained. The age-interactive model provided a better fit than the age-additive model. **c. Visualisation of the interaction effects from the age-interactive model**. Observed g-scores are plotted against within-person age (age*_dev_*), with individual trajectories coloured by mean-centered between-person age (age*_mean_*). The interaction indicates that the within-person age-related decline in cognitive functioning was steeper among older adults (lighter blue) than among younger adults (darker blue).

### Performance of predictive models based on individual neuroimaging phenotypes

We first assessed the performance of five machine-learning algorithms in predicting the observed g-score from each neuroimaging phenotype. Supplementary Table 3 and Supplementary Figure 1 summarise the predictive performance, measured as the Pearson correlation (r) between predicted and observed g-scores, for five machine-learning algorithms: Random Forest, Elastic Net, XGBoost, Partial Least Squares (PLS), and Kernel Ridge Regression (KRR). Predictive performance varied considerably by modality: diffusion-weighted imaging (DWI) and structural MRI (sMRI) demonstrated the strongest performance, followed by functional connectivity (FC) and task-based functional MRI (task-fMRI), with arterial spin labelling (ASL) consistently the weakest predictor.

### Diffusion-Weighted Imaging (DWI)

DWI-based models outperformed all other neuroimaging modalities, with Pearson correlations ranging from 0.51 to 0.68 across algorithms. While algorithm choice had a modest impact, DWI-derived connectomic features (r = 0.63-0.68) consistently provided a more robust basis for predicting cognitive variation than tractometry features, such as fractional anisotropy (r = 0.52-0.56) and mean diffusivity (r = 0.60-0.61).

### Structural MRI (sMRI)

sMRI morphometric measures also demonstrated strong predictive performance, with correlations ranging from 0.4 to 0.69. Across all algorithms, subcortical volumes consistently provided the highest correlations, peaking at r = 0.69 under Random Forest. Global brain variables and cortical thickness also showed robust predictive power (r = 0.63-0.69), while grey-matter volume performed moderately (r = 0.54), and cortical surface area was consistently the weakest structural predictor (r = 0.37-0.39).

### Functional connectivity (FC)

FC-based models, including resting-state and task-based connectivity networks, showed moderate predictive performance. Resting-state connectivity correlations ranged from 0.47 to 0.55 across algorithms. In contrast, task-based functional connectivity, defined here as connectivity calculated on residual BOLD fluctuations after removing task-evoked responses, showed improved performance (r = 0.57-0.68).

### Task-fMRI

Across all algorithms, task-fMRI models provided moderate predictive power. The correlation between predicted and observed cognitive scores ranged between 0.3 and 0.54, depending on algorithms and tasks. Among the four task contrasts, the scene-viewing condition gave the strongest predictions, peaking at r = 0.54 with Elastic Net. The semantic difficulty contrast from the Words task followed closely, achieving r = 0.53 under Random Forest. Algorithm choice again had only a secondary influence, compared to differences between contrasts.

### Arterial Spin Labeling (ASL)

ASL-derived cerebral blood flow (CBF) measures consistently delivered the weakest predictive performance across all tested algorithms, with correlations ranging from approximately 0.01 to 0.36. The best-performing regional CBF maps were grey-matter-corrected, achieving a maximum correlation of 0.36 with Random Forests using HCP parcellations.

### Comparison across algorithms

Across the 37 feature sets, algorithm choice had only a modest impact on accuracy: the median difference in correlation between the best-and worst-performing models was just 0.05 (IQR = 0.03-0.08). Nevertheless, Elastic Net either tied for or held the leading position in four out of five neuroimaging modalities. Specifically, Elastic Net matched Random Forest for sMRI (maximum r = 0.69), tied with XGBoost, PLS, and KRR for DWI (r = 0.68), was among the highest-performing methods alongside PLS and KRR for FC (r = 0.68), and uniquely led for task-fMRI (r = 0.54). For ASL, Random Forest and XGBoost jointly achieved the highest correlation (r = 0.36).

### Predictive performance of stacked models

Next, we assessed whether combining phenotypes via stacking could improve predictive accuracy. As summarised in Figure 3 and Supplementary Table 4, this approach consistently enhanced performance over the best individual phenotype models, increasing the mean Pearson correlation by 0.02-0.12 and reducing the Mean Absolute Error (MAE) by 0.02–0.13. Combining all 37 neuroimaging phenotypes into a multimodal stacking model yielded the strongest overall predictive performance, achieving a mean Pearson correlation of 0.73 and an R² of 0.51 across the five outer cross-validation folds.

**Figure 3.**
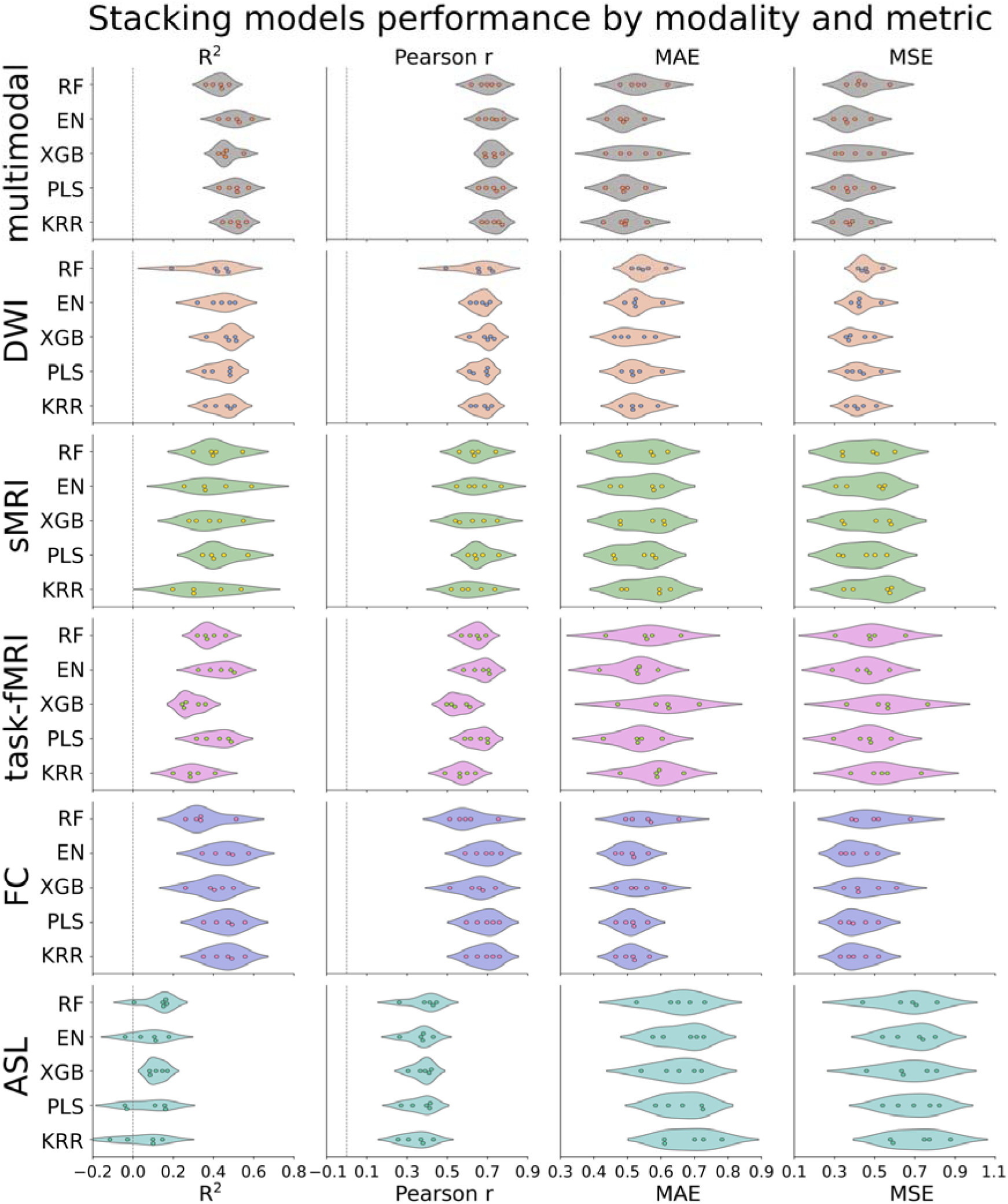
Predictive performance of stacked models, comparing outcomes across neuroimaging modalities and evaluation metrics. Violin plots show performance across five outer cross-validation folds for each stacked model. Rows mark the neuroimaging modality in the stacking (multimodal, DWI, sMRI, task-fMRI, FC, ASL). Columns display the evaluation metrics: coefficient of determination (R²), Pearson correlation coefficient (r), mean absolute error (MAE) and mean squared error (MSE). Points are fold values, the violin width shows their distribution. Algorithms on the ordinate are (from top): Random Forest (RF), Elastic Net (EN), XGBoost (XGB), Partial Least Squares (PLS) and Kernel Ridge Regression (KRR). Detailed results are available in the Supplementary Table 4. *Abbreviations*: DWI – diffusion-weighted imaging; sMRI – structural magnetic resonance imaging; task-fMRI – task-based functional magnetic resonance imaging; FC – functional connectivity; ASL – arterial spin labelling.

Stacking phenotypes within each modality revealed a similar, though slightly altered, performance hierarchy. DWI remained the strongest single modality (r = 0.69, R² = 0.47), but the performance of FC was substantially enhanced, placing it nearly on par with DWI (r = 0.69, R² = 0.46). These were followed by sMRI and task-fMRI, which showed identical predictive performance (r = 0.66, R² = 0.43). Notably, sMRI was the only modality for which stacking provided no performance gain over its best individual phenotype. Finally, ASL again showed the weakest performance (r = 0.39, R² = 0.13).

### Feature importance

To identify which neuroimaging features were most predictive, we examined feature importances for the best-performing model in each modality (Figure 4; Supplementary Tables 5-25). We estimated the contribution of each feature by calculating the Pearson correlation between its scaled values and the corresponding predicted g-scores ^40^. The resulting correlation coefficient thus indicates how strongly each feature contributed to the model’s predictions of cognitive functioning.

**Figure 4.**
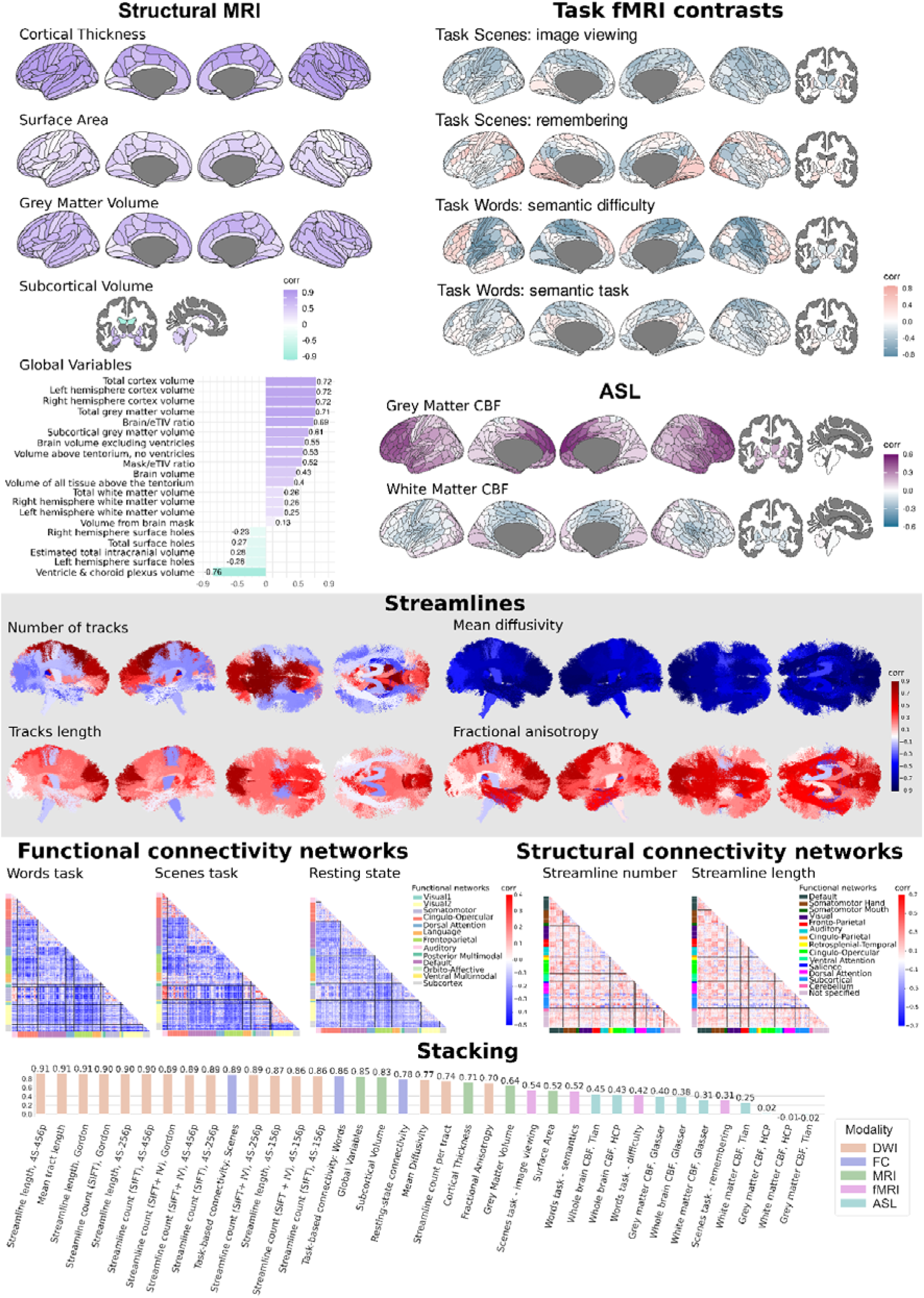
Feature Importance of the top-performing algorithms for each neuroimaging phenotype and multimodal stacking. A higher absolute correlation value indicates a higher contribution to the prediction. The detailed versions of the feature importance for each set of neuroimaging features can be found in Supplementary Tables 5-25. Abbreviations: DWI – diffusion-weighted imaging; sMRI – structural magnetic resonance imaging; task-fMRI – task-based functional magnetic resonance imaging; FC – functional connectivity; ASL – arterial spin labelling; CBF – cerebral blood flow; SIFT2 – Spherical-deconvolution Informed Filtering of Tractograms; 4S-***p (e.g., 4S-156p) – Schaefer atlas with *** parcels; HCP – Human Connectome Project parcellation that includes a combination of Glasser and ASEG atlases.

An analysis of feature importance for the final multimodal stacking model revealed that its predictive performance was driven by measures of structural connectivity derived from DWI. Features related to streamline length and count consistently ranked as the most influential, with nine of the top ten features belonging to the DWI modality and exhibiting correlations with the final prediction above 0.89. Following these, task-based functional connectivity and sMRI metrics, particularly global brain variables and subcortical volumes, also demonstrated substantial importance (r > 0.83). In contrast, task-evoked activation from task-fMRI and especially ASL-derived cerebral blood flow were far less informative. Several ASL features showed near-zero or negative correlations, indicating they contributed negligible predictive value to the final stacked model.

### Variance Decomposition

To assess the extent to which MRI markers from different predictive models explain within-and between-person variation in cognitive functioning (g-score), we performed variance decomposition on the neuroimaging-only models ^38^. These mixed-effects models simultaneously included two fixed effects: (1) a between-person MRI marker, defined as the individual’s average predicted g-score across waves, and (2) a within-person MRI marker, defined as the deviation of an individual’s MRI marker from their own mean across waves.

Figure 5 presents the beta estimates from the neuroimaging-only models ^38^. Neuroimaging phenotypes from most modalities, except ASL, yielded statistically significant β values (p <.05) for both between-person and within-person MRI markers. The magnitude of the between-person β was consistently greater than that of the within-person β. For example, the multimodal stacking model, which integrates information across modalities, produced β = 0.77 (SE = 0.04, p <.001) for the between-person MRI marker and β = 0.3 (SE = 0.04, p <.001) for the within-person MRI marker, indicating a stronger unique contribution from between-person variation (see also Figure 7a for model summary). Detailed results for mixed-effects models are available in the Supplementary Tables 26-32.

**Figure 5.**
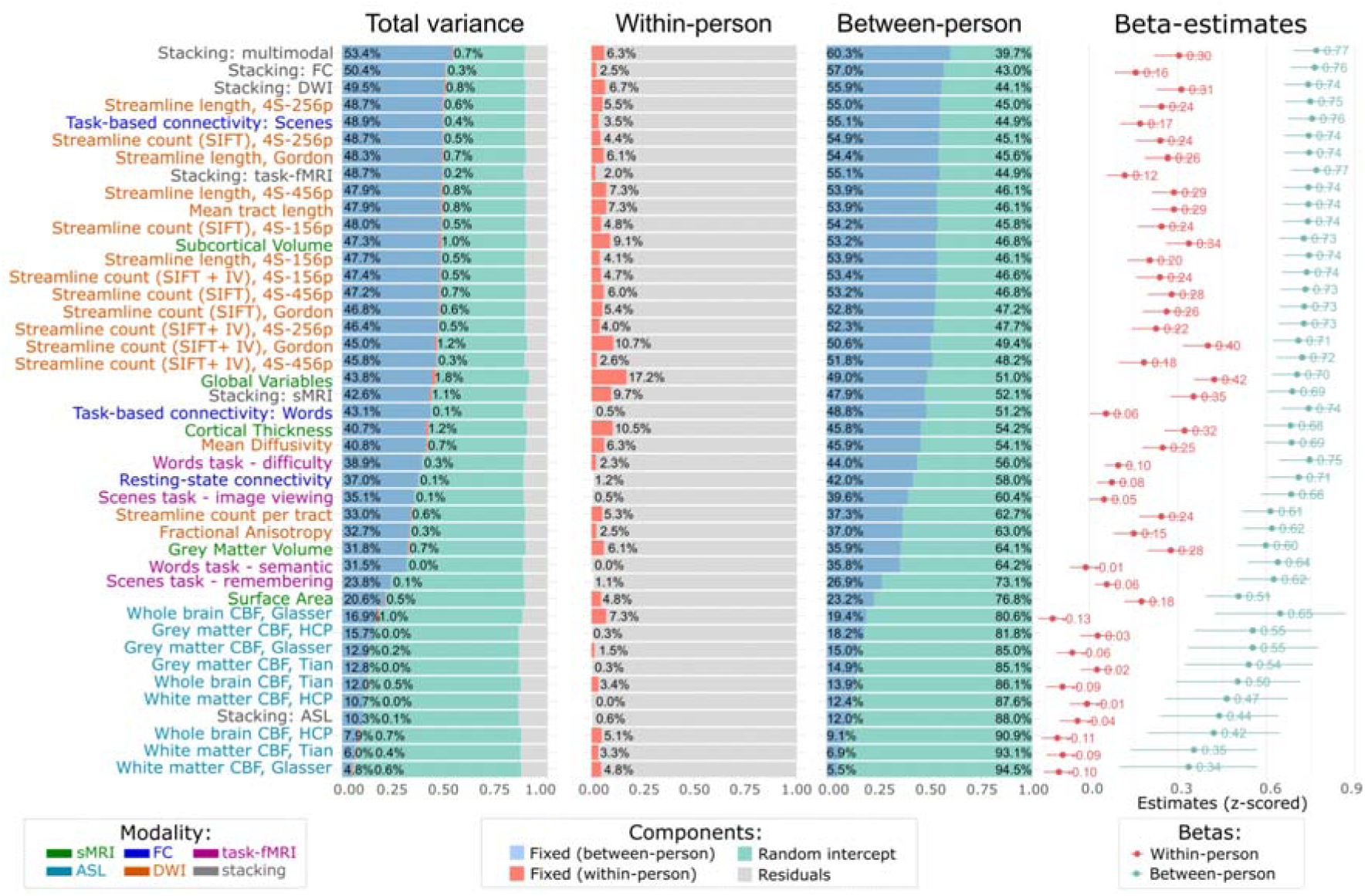
The variance-decomposition and corresponding. β **estimates and their 95%CI for the neuroimaging-only model.** The figure summarises how much of the total, within-person and between-person variance in cognitive functioning (g-score) is explained by MRI markers from each modality, along with their fixed-effect estimates. The leftmost panel shows, for each given individual neuroimaging phenotype or stacking, the proportion of total variance in observed g-scores explained by fixed and random effects. Rows ordered by total fixed-effect share. The central panels isolate the fixed-effect share into between-person (mean predicted g-score across waves) and within-person (deviation from each person’s mean) components, expressed as a percentage of total variance. The rightmost panel plots z-scored beta coefficients with 95 % confidence intervals for within-person and between-person effects from the mixed model that includes only MRI predictors. *Abbreviations*: DWI – diffusion-weighted imaging; sMRI – structural magnetic resonance imaging; task-fMRI – task-based functional magnetic resonance imaging; FC – functional connectivity; ASL – arterial spin labelling; CBF – cerebral blood flow; SIFT2 – Spherical-deconvolution Informed Filtering of Tractograms; 4S-***p (e.g., 4S-156p) – Schaefer atlas with *** parcels; HCP – Human Connectome Project parcellation that includes a combination of Glasser and ASEG atlases.

Figure 5 also reports variance decomposition, partitioning the total variance in observed cognitive functioning into within-person and between-person components. For the within-person component, the fixed effect of the within-person MRI markers explained up to 17.2% of this variance (global variables from sMRI). Across stacking models, the proportion of within-person variance captured by MRI markers ranged from 0.6% (ASL) to 2% (task-fMRI), 2.5% (FC), 6.7% (DWI), 6.3% (multimodal), and 9.7% (sMRI).

In contrast, for the between-person component, the fixed effect of the between-person MRI markers explained up to 60.3% of this component (multimodal stacking). Across modality-specific stacking models, the proportion of between-person variance captured by MRI markers ranged from 12% (ASL) to 47.9% (sMRI), 55.1% (task-fMRI), 55.9% (DWI) and 57% (FC). Detailed results for variance decomposition are available in the Supplementary Table 33.

Figure 6 visualises the capacity of the MRI-based stacking models in capturing between-person versus within-person variability in cognitive functioning. For the between-person variability, we computed the mean of both observed and predicted g-score across waves and examined their correlations. These correlations were all positive and statistically significant (p<.001), ranging from r=.49 (ASL) to r≥.70 for other modalities. For within-person variability, we computed the deviations of observed and predicted g-scores from their respective means across waves and examined their correlations. Apart from ASL, these correlations were statistically significant (p<.001), ranging from r=.18 (task-fMRI) to r=.39 (sMRI).

**Figure 6.**
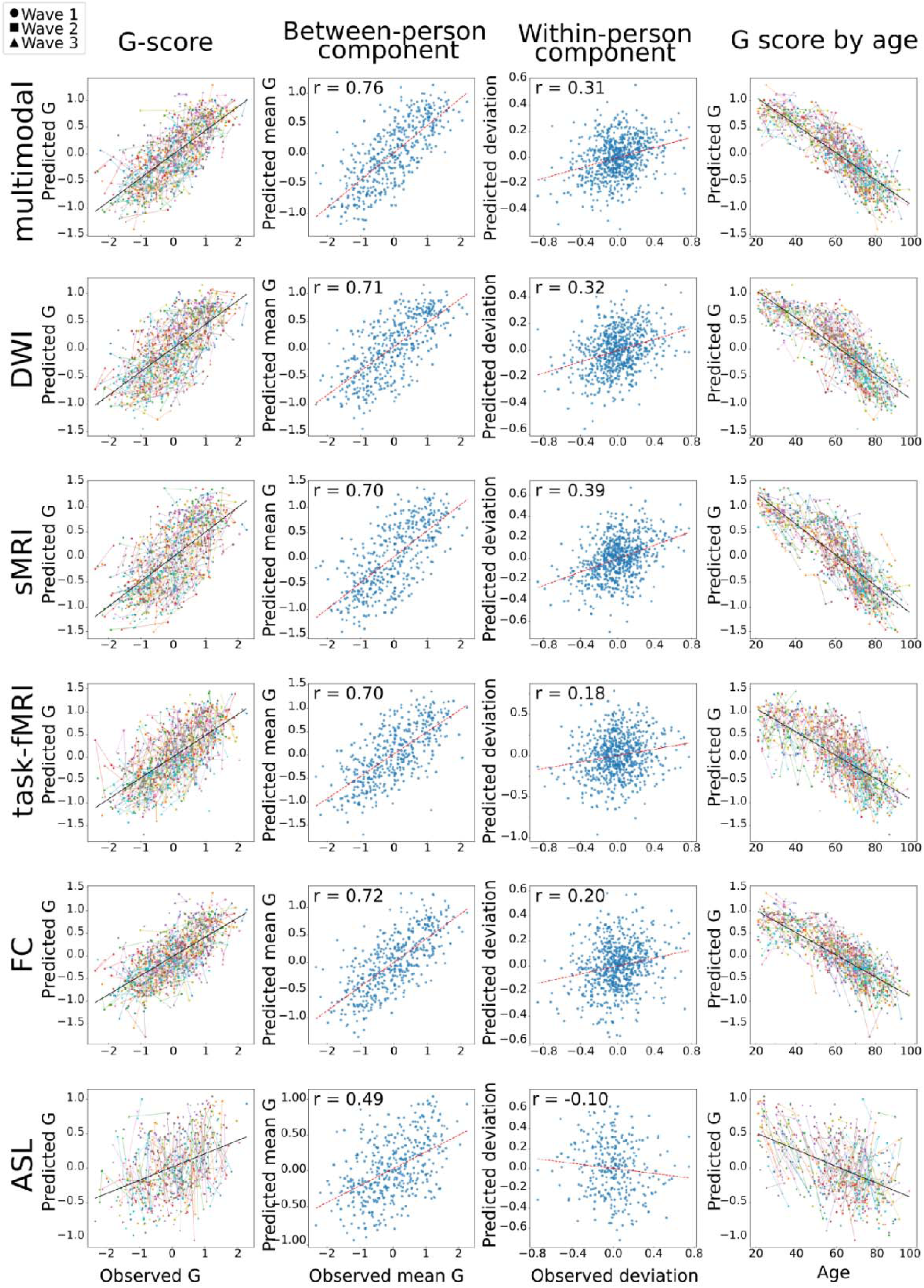
Prediction of g-score across MRI modalities, separating between-and within-person variability, and relation with age. Rows show the multimodal stacking and modality specific stacking for DWI, sMRI, task-fMRI, FC, and ASL. For each stacking model four panels are shown. Column 1: Each point represents the correspondence between an observed and predicted g-score from one participant, with solid lines connecting repeated observations from the same individual over up to three waves. Colours distinguish individuals only to facilitate visual tracking, and marker shapes indicate the wave. Black solid line depicts correlation between observed and predicted values. Column 2: Each point plots a participant’s mean predicted g-score against their mean observed g-score across all available waves, isolating stable between-person differences, to indicate how well the model captures differences between people. The red line and the accompanying r (Pearson coefficient) show the correlation between observed and predicted values. Column 3: Each point shows the within-person deviation of the predicted g-score (wave value minus that participant’s mean) against the corresponding deviation in the observed g-score to indicate how well the model tracks longitudinal change within individuals. The red line and the accompanying r (Pearson coefficient) show the correlation between observed and predicted values. Column 4: Predicted g-scores are plotted against chronological age. Points and coloured lines follow the same individual and wave coding as in Column 1, so trajectories across up to three waves can be tracked. The black solid line depicts the overall age trend captured by the model. *Abbreviations*: DWI – diffusion-weighted imaging; sMRI – structural magnetic resonance imaging; task-fMRI – task-based functional magnetic resonance imaging; FC – functional connectivity; ASL – arterial spin labelling.

**Figure 7.**
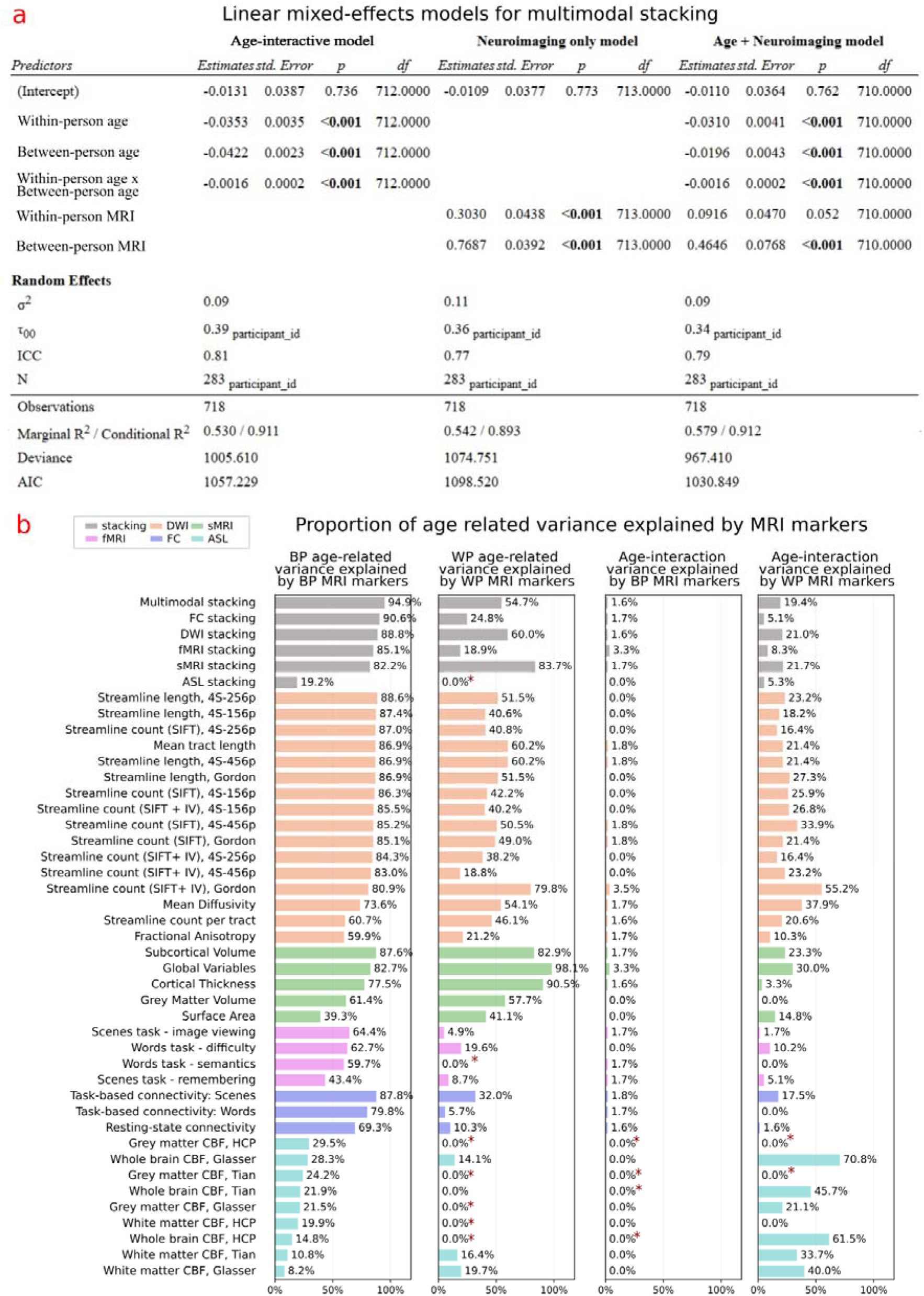
Linear mixed-effects models and commonality analysis of variance in cognitive functioning. The figure illustrates the extent to which machine learning predictors trained on neuroimaging features capture the effect of age on cognitive functioning (g-score), and how much age-related variance they capture, while separating within-person from between-person sources of variance. a. The table reports results for three linear mixed-effects models for multimodal stacking: an age-interactive model that enters the age terms as predictors of cognitive functioning (g-score); a neuroimaging-only model that enters predicted cognitive scores from multimodal stacking; and a joint model that includes both age and neuroimaging as predictors. All models use repeated observations nested within participants, allowing separate estimation of within-person and between-person effects. b. Bar plots from the commonality analysis of the age-plus-neuroimaging models for the multimodal and modality specific stacking models. The four panels show the proportion of variance shared between age terms and MRI markers. Colours indicate the stacking model and the modality specific models derived from DWI, sMRI, task-fMRI, FC, and ASL. Note: * indicates negative fractions, which can arise due to sampling noise or multicollinearity, were set to zero. *Abbreviations*: DWI – diffusion-weighted imaging; sMRI – structural magnetic resonance imaging; task-fMRI – task-based functional magnetic resonance imaging; FC – functional connectivity; ASL – arterial spin labelling; CBF – cerebral blood flow; SIFT2 – Spherical-deconvolution Informed Filtering of Tractograms; 4S-***p (e.g., 4S-156p) – Schaefer atlas with *** parcels; HCP – Human Connectome Project parcellation that includes a combination of Glasser and ASEG atlases.

### Commonality analysis

To assess the extent to which MRI markers explained the relationship between cognitive functioning and age, we conducted a commonality analysis^41,42^ on the age-plus-neuroimaging models. These linear mixed-effects models included five fixed effects: age*_dev_*, age*_mean_*, the age*_dev_*: age*_mean_* interaction, *g_pred_dev,_* and *g_pred_mean_*. Table 1 and Figure 7 summarise the results for the multimodal stacking markers and present commonality analyses across all MRI modalities. Below, we quantify the degree to which the MRI marker accounted for the associations between cognitive functioning and the three age-related variables, using the multimodal MRI marker as an example (for other MRI markers, see Supplementary Table 34).

**Table 1.**
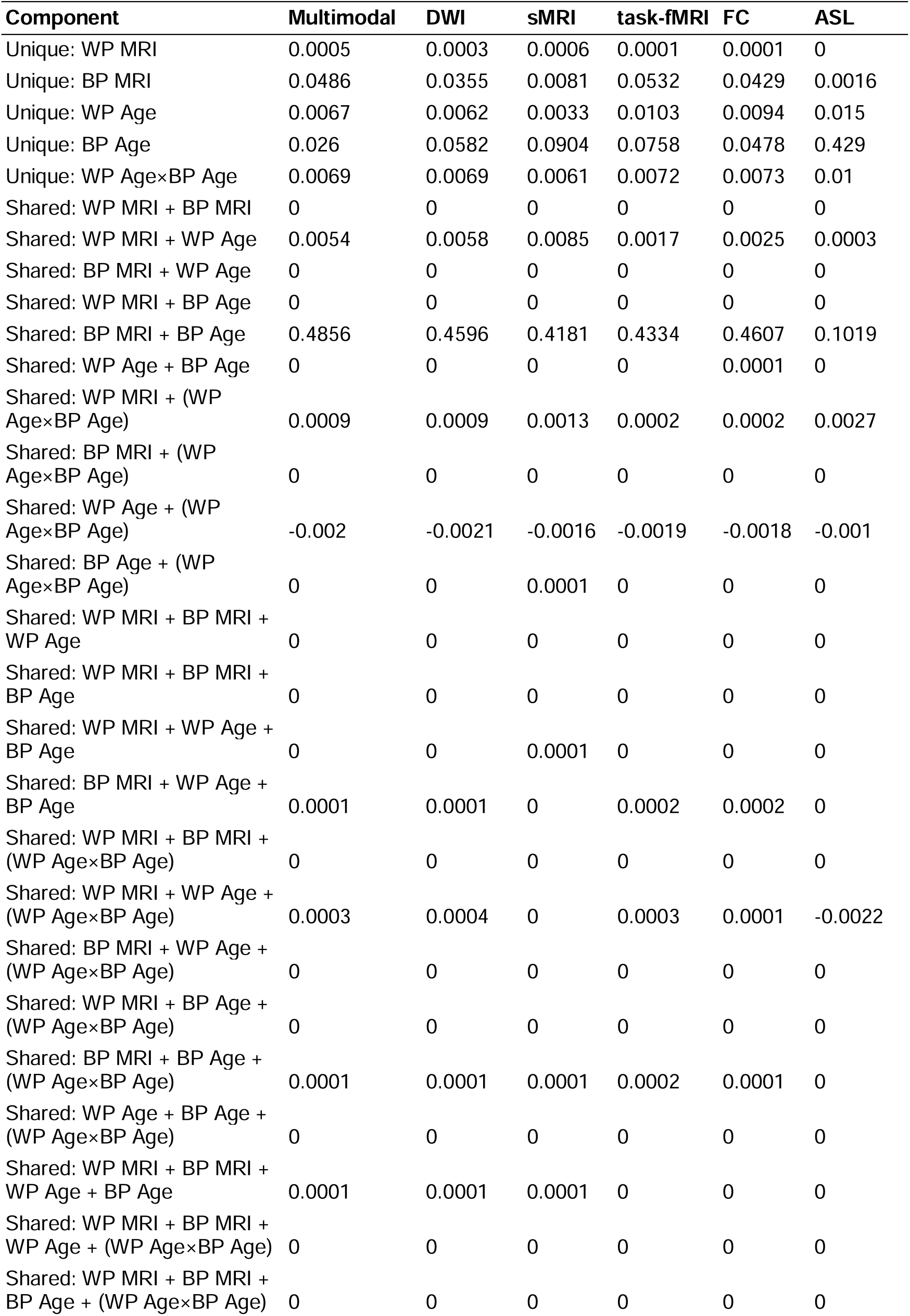
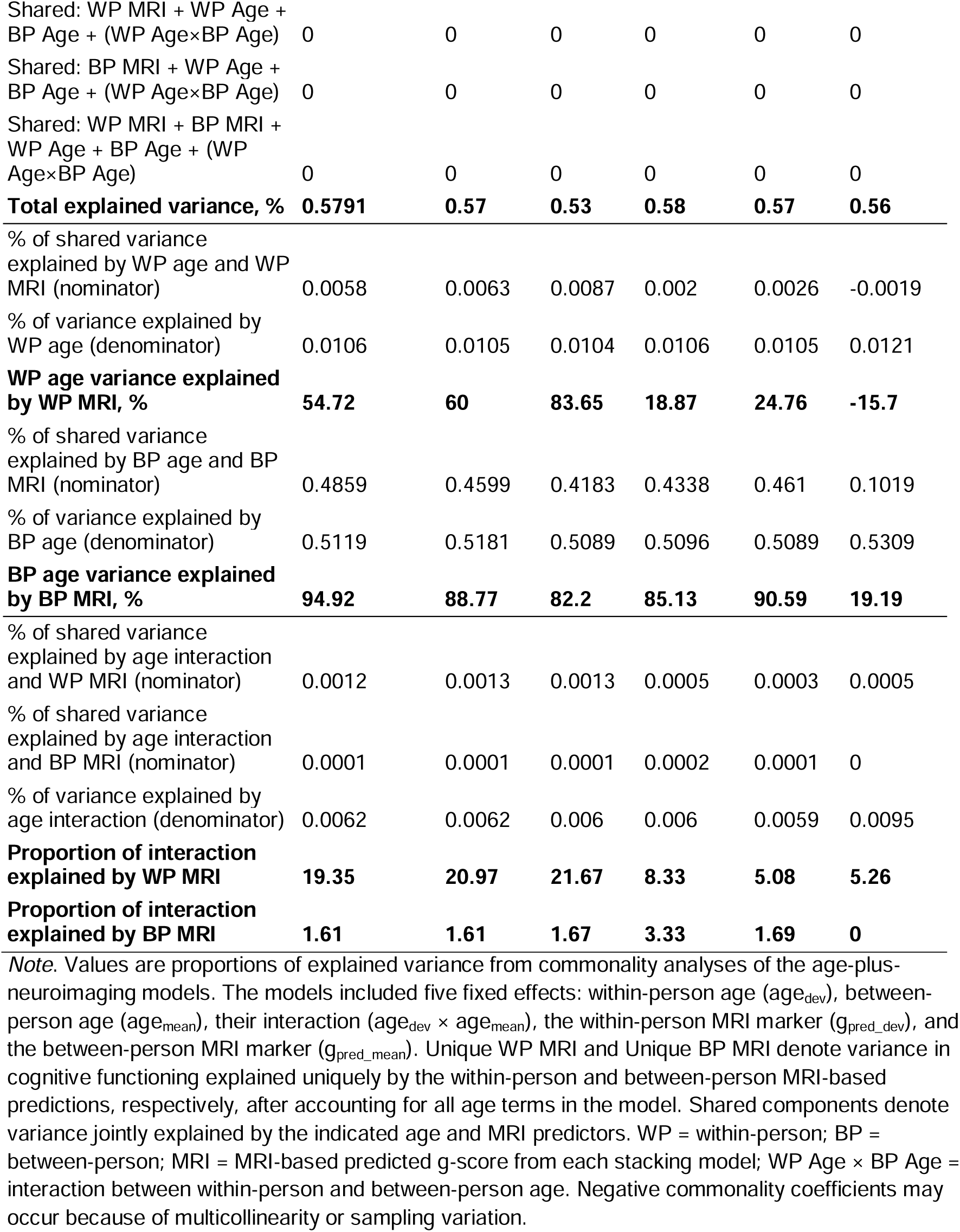
Variance partitioning of age-related effects explained by MRI markers, decomposed into within-person and between-person components of stacking models.

| Component | Multimodal | DWI | sMRI | task-fMRI | FC | ASL |
| --- | --- | --- | --- | --- | --- | --- |
| Unique: WP MRI | 0.0005 | 0.0003 | 0.0006 | 0.0001 | 0.0001 | 0 |
| Unique: BP MRI | 0.0486 | 0.0355 | 0.0081 | 0.0532 | 0.0429 | 0.0016 |
| Unique: WP Age | 0.0067 | 0.0062 | 0.0033 | 0.0103 | 0.0094 | 0.015 |
| Unique: BP Age | 0.026 | 0.0582 | 0.0904 | 0.0758 | 0.0478 | 0.429 |
| Unique: WP AgexBP Age | 0.0069 | 0.0069 | 0.0061 | 0.0072 | 0.0073 | 0.01 |
| Shared: WP MRI + BP MRI | 0 | 0 | 0 | 0 | 0 | 0 |
| Shared: WP MRI + WP Age | 0.0054 | 0.0058 | 0.0085 | 0.0017 | 0.0025 | 0.0003 |
| Shared: BP MRI + WP Age | 0 | 0 | 0 | 0 | 0 | 0 |
| Shared: WP MRI + BP Age | 0 | 0 | 0 | 0 | 0 | 0 |
| Shared: BP MRI + BP Age | 0.4856 | 0.4596 | 0.4181 | 0.4334 | 0.4607 | 0.1019 |
| Shared: WP Age + BP Age | 0 | 0 | 0 | 0 | 0.0001 | 0 |
| Shared: WP MRI + (WP AgexBP Age) | 0.0009 | 0.0009 | 0.0013 | 0.0002 | 0.0002 | 0.0027 |
| Shared: BP MRI + (WP AgexBP Age) | 0 | 0 | 0 | 0 | 0 | 0 |
| Shared: WP Age + (WP AgexBP Age) | -0.002 | -0.0021 | -0.0016 | -0.0019 | -0.0018 | -0.001 |
| Shared: BP Age + (WP AgexBP Age) | 0 | 0 | 0.0001 | 0 | 0 | 0 |
| Shared: WP MRI + BP MRI + WP Age | 0 | 0 | 0 | 0 | 0 | 0 |
| Shared: WP MRI + BP MRI + BP Age | 0 | 0 | 0 | 0 | 0 | 0 |
| Shared: WP MRI + WP Age + BP Age | 0 | 0 | 0.0001 | 0 | 0 | 0 |
| Shared: BP MRI + WP Age + BP Age | 0.0001 | 0.0001 | 0 | 0.0002 | 0.0002 | 0 |
| Shared: WP MRI + BP MRI + (WP AgexBP Age) | 0 | 0 | 0 | 0 | 0 | 0 |
| Shared: WP MRI + WP Age + (WP AgexBP Age) | 0.0003 | 0.0004 | 0 | 0.0003 | 0.0001 | -0.0022 |
| Shared: BP MRI + WP Age + (WP AgexBP Age) | 0 | 0 | 0 | 0 | 0 | 0 |
| Shared: WP MRI + BP Age + (WP AgexBP Age) | 0 | 0 | 0 | 0 | 0 | 0 |
| Shared: BP MRI + BP Age + (WP AgexBP Age) | 0.0001 | 0.0001 | 0.0001 | 0.0002 | 0.0001 | 0 |
| Shared: WP Age + BP Age + (WP AgexBP Age) | 0 | 0 | 0 | 0 | 0 | 0 |
| Shared: WP MRI + BP MRI + WP Age + BP Age | 0.0001 | 0.0001 | 0.0001 | 0 | 0 | 0 |
| Shared: WP MRI + BP MRI + WP Age + (WP AgexBP Age) | 0 | 0 | 0 | 0 | 0 | 0 |
| Shared: WP MRI + BP MRI + BP Age + (WP AgexBP Age) | 0 | 0 | 0 | 0 | 0 | 0 |
| Shared: WP MRI + WP Age + BP Age + (WP Age×BP Age) | 0 | 0 | 0 | 0 | 0 | 0 |
| Shared: BP MRI + WP Age + BP Age + (WP Age×BP Age) | 0 | 0 | 0 | 0 | 0 | 0 |
| Shared: WP MRI + BP MRI + WP Age + BP Age + (WP Age×BP Age) | 0 | 0 | 0 | 0 | 0 | 0 |
| **Total explained variance, %** | **0.5791** | **0.57** | **0.53** | **0.58** | **0.57** | **0.56** |
| % of shared variance explained by WP age and WP MRI (nominator) | 0.0058 | 0.0063 | 0.0087 | 0.002 | 0.0026 | -0.0019 |
| % of variance explained by WP age (denominator) | 0.0106 | 0.0105 | 0.0104 | 0.0106 | 0.0105 | 0.0121 |
| **WP age variance explained by WP MRI, %** | **54.72** | **60** | **83.65** | **18.87** | **24.76** | **-15.7** |
| % of shared variance explained by BP age and BP MRI (nominator) | 0.4859 | 0.4599 | 0.4183 | 0.4338 | 0.461 | 0.1019 |
| % of variance explained by BP age (denominator) | 0.5119 | 0.5181 | 0.5089 | 0.5096 | 0.5089 | 0.5309 |
| **BP age variance explained by BP MRI, %** | **94.92** | **88.77** | **82.2** | **85.13** | **90.59** | **19.19** |
| % of shared variance explained by age interaction and WP MRI (nominator) | 0.0012 | 0.0013 | 0.0013 | 0.0005 | 0.0003 | 0.0005 |
| % of shared variance explained by age interaction and BP MRI (nominator) | 0.0001 | 0.0001 | 0.0001 | 0.0002 | 0.0001 | 0 |
| % of variance explained by age interaction (denominator) | 0.0062 | 0.0062 | 0.006 | 0.006 | 0.0059 | 0.0095 |
| **Proportion of interaction explained by WP MRI** | **19.35** | **20.97** | **21.67** | **8.33** | **5.08** | **5.26** |
| **Proportion of interaction explained by BP MRI** | **1.61** | **1.61** | **1.67** | **3.33** | **1.69** | **0** |

*Note.* Values are proportions of explained variance from commonality analyses of the age-plus-neuroimaging models. The models included five fixed effects: within-person age ( $age_{dev}$ ), between-person age ( $age_{mean}$ ), their interaction ( $age_{dev} \times age_{mean}$ ), the within-person MRI marker ( $g_{pred\_dev}$ ), and the between-person MRI marker ( $g_{pred\_mean}$ ). Unique WP MRI and Unique BP MRI denote variance in cognitive functioning explained uniquely by the within-person and between-person MRI-based predictions, respectively, after accounting for all age terms in the model. Shared components denote variance jointly explained by the indicated age and MRI predictors. WP = within-person; BP = between-person; MRI = MRI-based predicted g-score from each stacking model; WP Age × BP Age = interaction between within-person and between-person age. Negative commonality coefficients may occur because of multicollinearity or sampling variation.

In addition to the shared age-related components, the commonality analysis identified unique effects of the MRI-based predictors. The unique effect of g_pred_mean_ represents between-person variance in cognitive functioning explained by the MRI-based prediction after accounting for age_mean_, age_dev_, and their interaction, that is, age-independent differences between individuals captured by the MRI marker. The unique effect of g_pred_dev_ represents within-person variance in cognitive functioning explained by within-person fluctuations in the MRI-based prediction after accounting for the age terms, that is, age-independent longitudinal covariation between MRI-based prediction and cognitive functioning within individuals.

The main effect of age*_dev_*, representing within-person age-related change at the mean of age*_mean_*, explained 1.06% of the total variance in observed g-scores. The shared variance between age*_dev_* and *g_pred_dev_* from the multimodal MRI marker, aggregated across all combinations of fixed-effect variables, was 0.58%. Thus, the multimodal MRI marker accounted for 0.58% ÷ 1.06% = 54.72% of the within-person age-related variance.

The main effect of age*_mean_*, representing between-person age differences at the mean of age*_dev_*, explained 51.19% of the total variance in observed g-scores. The shared variance between age*_mean_* and *g_pred_mean_* across all combinations of fixed-effect variables was 48.59%. Therefore, the multimodal MRI marker captured 48.59% ÷ 51.19% = 94.92% of the between-person age-related variance.

The interaction between age*_dev_* and age*_mean_*, reflecting how within-person age effects vary as a function of between-person age, accounted for 0.62% of the total variance in observed g-scores. The interaction shared 0.12% of its variance with *g_pred_dev_* and 0.01% with *g_pred_mean_*. Accordingly, the multimodal MRI marker explained 0.12% ÷ 0.62% = 19.35% (via *g_pred_dev_*) and 0.01% ÷ 0.62% = 1.61% (via *g_pred_mean_*) of the age-interaction variance.

## Discussion

This study benchmarked the ability of MRI markers from five modalities to capture within-and between-person variation in cognitive functioning. We pursued three aims. First, we compared predictive performance across neuroimaging phenotypes and modalities. The multimodal stacking, which integrates features from all five modalities, achieved the highest accuracy, followed by DWI, FC, and sMRI. Second, we quantified each MRI marker’s capacity to explain within-versus between-person variation. Variance decomposition revealed an asymmetry: MRI markers were highly effective for predicting between-person differences but only modest, though significant, at tracking within-person longitudinal changes. Third, we examined how well these markers captured the cognition–age relationship. In this dataset, the cognition–age association was substantial between persons but considerably weaker within persons; nonetheless, most MRI markers were able to capture both components of this relationship.

Our first objective was to determine the predictive performance of various MRI markers for cognitive functioning. In line with a growing body of research ^18–22^, the multimodal stacking model, which combined information from all 37 neuroimaging phenotypes, yielded the highest predictive accuracy. Stacking phenotypes within each modality also consistently improved performance over individual measures, supporting the view that different imaging phenotypes provide complementary, while overlapping, biological information relevant to cognition.

Predictive accuracy was highly dependent on the MRI modality, and even within a single modality, performance varied substantially based on the quantification method and the brain atlas used for parcellation. Diffusion features, especially structural connectivity measures, were the main drivers of the multimodal stacking and delivered strong single modality performance. This aligns with correlational studies that consistently link the integrity of white matter tracts to cognitive abilities that decline with age, such as processing speed and executive function ^43^. While individual functional connectivity (FC) measures were moderately predictive, stacking them across tasks and resting state elevated performance, reaching parity with DWI predictions. Previous research consistently found FC among the strongest predictors of cognition ^10,20,44,45^. Echoing earlier findings ^46–48^, task-derived FC outperformed resting-state FC in our analysis. Structural MRI followed closely and, interestingly, some individual morphometric measures, such as subcortical volume and global variables, outperformed the sMRI stacking. Meta-analyses confirm a strong correlation between brain volume and intelligence ^49,50^, and out-of-sample studies often report robust predictions from global and regional brain volumes ^15,51^. Task fMRI contrasts performed moderately, though stacking significantly enhanced predictive accuracy (11.5% up to 25.3% boost in explained variance). Consistent with Tetereva and colleagues ^20^, the relatively undemanding tasks available in the Dallas Lifespan Brain Study (DLBS) yielded predictive performance comparable to that of low-cognitive-demanding paradigms reported previously.

Although prior research has highlighted ASL as a potentially promising predictor of cognitive change ^52–54^, ASL-derived features were consistently the weakest predictors in our models, lagging significantly behind other modalities. We attribute this discrepancy to inherently low signal-to-noise ratio (SNR), which arises because the perfusion signal from labelled arterial blood is a small fraction (often <1%) of the total tissue signal ^55,56^. This low SNR leads to challenges in image quality, quantification accuracy, and clinical reliability, especially in populations with low blood flow or in high-resolution imaging ^57^. The quality evaluation index in our preprocessing flagged the available ASL data as suboptimal. We therefore refrain from drawing firm conclusions about its broader utility of ASL for predicting cognition.

Our second objective was to evaluate the capacity of each MRI marker to explain within-versus between-person variation in cognitive functioning. Statistically, most markers across modalities, except ASL, significantly captured both sources of variation, but the variance explained was highly asymmetric. Between-person markers accounted for a much larger proportion of cognitive variance than within-person markers. For example, the multimodal stacking model explained 60.3% (r =.76) of the between-person variance but only 6.3% (r =.31) of the within-person variance. We interpret this pattern in the context of the Dallas Lifespan Brain Study (DLBS), which excluded individuals with major psychiatric or neurological disorders. Because the sample comprised generally healthy adults, cognitive functioning showed little change over the 5–10-year period, as reflected in the high intraclass correlation coefficient (ICC) for the g-score. An ICC of 0.88 indicates that 88% of the total variance in cognitive functioning occurred between individuals and only 12% within individuals (see Supplementary Table 35 for the ICC of the MRI-based predicted g-scores). Future work examining samples with more pronounced cognitive decline, such as some forms of dementia, may reveal different levels of sensitivity for MRI markers in capturing within-person variation in cognitive functioning.

The between-person component was largest for models that stacked diffusion and functional connectivity features. At the phenotype level, this signal was driven by structural connectivity metrics, such as streamline count and length, and key morphometric measures, including subcortical volumes and global brain variables. For the within-person component, sMRI-based phenotypes showed the most sensitivity to longitudinal change (up to 17.2% from the sMRI global variables). ASL-based predictors were negligible at both levels.

The variance decomposition across modalities offers a useful guide for future research. For studies aiming to explain stable, between-person cognitive differences, especially when resources are constrained, DWI and stacked FC models represent robust and efficient choices. However, for research focused on tracking longitudinal within-person change, sMRI features showed the most promise, though the overall subtlety of this signal underscores the considerable challenge of predicting individual cognitive trajectories, especially among healthy participants.

Our third objective was to examine the extent to which the MRI-based models captured age-related variance in cognitive functioning. We first quantified the relationship between age and cognitive functioning. The main effect of between-person differences in age (age*_mean_*) accounted for a substantially larger proportion of variance in cognitive functioning (51.15%) than either the main effect of within-person changes in age (age*_dev_*) (1.06%) or the interaction between the two age components (0.62%). This pattern likely reflects the healthy nature of the DLBS cohort. Despite this asymmetry, the MRI-based models were able to capture these variance components. For example, the between-person multimodal MRI marker accounted for most of the between-person age-related cognitive variance (94.92%). The within-person multimodal MRI marker captured more than half of the within-person age-related cognitive variance (54.72%) and also explained part of the age-interaction variance (19.35%). This accords with evidence that cognitive performance declines with age ^7^ and that MRI features are consistently predictors of age ^2,56^. ASL again diverged from this pattern, showing relatively low overlap with age, likely reflecting its overall poorer predictive performance.

The link between cognition and the brain is age-dependent and shaped by developmental, compensatory, and degenerative processes ^58,59^. Capturing age-related cognitive variation is therefore central to evaluating the translational utility of MRI markers. Because age accounts for a substantial portion of variance in cognitive functioning ^2,7,27^, MRI markers that recover this pattern are more likely to hold clinical value. In our data, both between-person and within-person MRI markers explained meaningful variance in cognition, and much of this variance overlapped with age.

This overlap validates that the between-person MRI markers capture aging-related biological processes, supporting their use in phenotyping and risk stratification. At the same time, although within-person MRI markers show potential for capturing within-person age-related cognitive change, confirming their sensitivity will require datasets with greater within-person cognitive variation.

The study has several limitations. First, neither of the two tasks in the DLBS fMRI paradigm was designed to be cognitively demanding. Task fMRI from paradigms, such as the n-back working memory task ^60^ and the episodic memory facename task ^61^, have produced some of the most robust MRI markers of cognition ^21^. By not including more demanding tasks, we may have missed condition-specific variance relevant to particular cognitive domains. Second, data collection preceded publication of the current ASL guidelines ^62^, so scanning parameters do not fully align with contemporary standards. This adds to the inherently low signal-to-noise ratio of ASL. The suboptimal ASL quality limits firm conclusions about the utility of this modality for predicting cognition, although more sophisticated methods may improve signal stability in future work ^63^. Next, sample characteristics may constrain generalisability. Participants in this study predominantly reflect a “WEIRD” demographic (Western, Educated, Industrialised, Rich, and Democratic; ^64^).

Education levels were relatively high, with 93.3% reporting at least some tertiary education and about one-fifth holding a postgraduate qualification. Self-reported race data indicated that 85% identified as White/Caucasian, with limited representation from other ethnic backgrounds. While socioeconomic status data were unavailable, the overall profile suggests a relatively privileged and homogeneous sample.

Extending these findings to more diverse populations remains an important direction for future research.

Taken together, our findings demonstrate that multimodal MRI can provide robust and generalisable predictions of cognitive functioning throughout adulthood. These predictors are particularly effective in capturing stable, trait-like differences between individuals and in accounting for age-related cognitive variation. For within-person differences, MRI modalities have the potential to track cognitive change as people age, but will need to be further investigated using data with higher within-person variability in cognitive functioning. Recognising this distinction is essential for translational applications, as it clarifies the current utility of MRI-based models for phenotyping and risk stratification, and highlights the methodological advances needed before such tools can be used reliably for individual-level monitoring over time.

## Methods

### Participants

We analysed data from the Dallas Lifespan Brain Study (DLBS), a publicly available longitudinal, multimodal neuroimaging dataset (DOI:10.18112/openneuro.ds004856.v1.0.2). The DLBS comprises three waves of data collection, each approximately 4-5 years apart, and includes participants aged 21 to 90 years. Ethics approval was obtained from the University of Texas Southwestern Medical Center Institutional Review Board (IRB #s: STU 072010-112, 072010-219, 092015-003).

The DLBS recruited right-handed individuals with at least a ninth-grade education, fluent in English, and a Mini-Mental State Examination (MMSE) score of ≥26 at baseline (Wave 1), with scores ≥22 accepted in subsequent waves. Participants were excluded if they had major psychiatric or neurological disorders, recent cancer treatment, a history of substance abuse, significant medical conditions, poor vision, excessive caffeine intake, elevated cholesterol or blood pressure, obesity (BMI ≥ 35), or contraindications to MRI scanning.

The DLBS provided structural magnetic resonance imaging (sMRI), task-based and resting-state functional MRI (fMRI), diffusion-weighted imaging (DWI), and arterial spin labeling (ASL), along with extensive cognitive and psychosocial assessments. Detailed scanning parameters and comprehensive demographic and individual difference measures (e.g., age, sex, education, ethnicity and race) were provided in the dataset documentation (DOI:10.18112/openneuro.ds004856.v1.0.2). For the present analyses, we included 450 participants (62.7% females, mean age_wave1_ = 57.8±17.6) whose cognitive scores were available across all 11 measurements used in the hierarchical modelling of general cognitive functioning. Supplementary tables 36 and 37 contain detailed information about the sample.

### Target: Cognitive score (g-score)

To estimate general cognitive functioning across cognitive measures, we derived a higher-order g-score using the exploratory-structural-equation-modelling within confirmatory-factor-analysis (ESEM-within-CFA) approach ^65^ with a five-fold cross-validation framework. The model was based on 11 cognitive measurements that were consistently administered across all waves and had minimal missing data, thereby maximizing the usable sample size. The cognitive tasks included Digit Comparison Task ^66^, WAIS-III Digit Symbol ^67^, WAIS-III Letter-Number Sequencing ^67^, CANTAB Spatial Working Memory ^68^, Hopkins Verbal Learning ^69^, Ravens Matrices ^70^, ETS Letter Sets ^71^, Cantab Stockings of Cambridge ^68^.

Cognitive test scores were z-standardised within each fold. We first evaluated data suitability for factor analysis using the Bartlett test of sphericity and the Kaiser–Meyer–Olkin (KMO) measure, both of which indicated acceptable factorability across folds. Bartlett’s test of sphericity was significant in all training folds (χ² > 4000, p < 0.001). KMO statistics ranged from 0.856 to 0.879, falling within the “meritorious” range ^72^.

We then applied principal axis exploratory factor analysis (EFA) with an oblique GeominQ rotation, retaining three factors based on parallel analysis and the interpretability of the factor structure across folds. The resulting factor structure consistently grouped cognitive tasks into three domains: working memory and reasoning, episodic memory, and processing speed. These three factors collectively explained 58% to 62% of the total variance.

Next, we constructed hierarchical models in which a higher-order general factor (g) loaded on the three lower-order factors. These models were estimated using the lavaan package ^39^, with maximum likelihood estimation. Fit indices indicated good model fit across folds: Root Mean Square Error of Approximation (RMSEA) ranged from 0.046 to 0.056, Comparative Fit Index (CFI) from 0.985 to 0.990, and Standardised Root Mean Square Residual (SRMR) from 0.018 to 0.023, Tucker–Lewis Index (TLI), consistently exceeded 0.96. Composite internal consistency of g-score was acceptable (mean ω = 0.74, range 0.72 to 0.76 across folds), indicating that the g-scores were measured with reasonable precision. Predicted latent scores for g and the three first-order factors were extracted separately for each fold’s training and test sets. Modelling details are provided in Supplementary Tables 1 and 2 and Figure 1.

### Features: Multimodal neuroimaging

We organized the neuroimaging data into 37 sets of features spanning task-based functional MRI (task-fMRI); functional connectivity (FC); structural MRI (sMRI); diffusion-weighted imaging (DWI); and arterial spin labeling (ASL). Acquisition parameters for each modality are described in detail in the original DLBS dataset documentation ^73^.

Feature sets were defined a priori to balance breadth across modalities with reproducibility. Rather than forcing a single atlas across all modalities, we used parcellations that are natively supported by each modality-specific preprocessing software containers (including QSIPrep ^74^, pyAFQ ^75^, fMRIPrep ^76^ and ASLPrep ^77^), because these align with how the underlying measures are estimated. Where a pipeline provided multiple parcellations or resolutions, we retained them to represent each modality at several spatial scales and parcellation strategies, rather than defaulting to a single atlas. For ASL, we retained only parcellations with adequate coverage and the least missing data.

Because our final multimodal model uses stacking, the inclusion of multiple parcellations or resolutions within a modality does not assume they are equally informative. Instead, the stacking model learns which phenotype-level predictions contribute most to out-of-sample prediction, effectively downweighting redundant or weak parcellations. We reported feature-importance analyses to show which underlying measures drive the strongest predictions.

### Diffusion-Weighted Imaging (DWI): 16 sets of features

We produced 16 feature sets from DWI data processed via single-shell three-tissue constrained spherical deconvolution and anatomically constrained probabilistic tractography. Preprocessing was performed using QSIPrep 1.0.1 ^74^. Supplementary Methods provide full preprocessing details.

Structural connectivity analyses yielded 12 feature sets. Whole-brain tractograms were parcellated with four atlases: three Schaefer-based ^78^ variants with 156, 256 and 456 regions, plus the 333-region Gordon atlas ^79^. For each atlas, we derived three connectivity matrices per participant: (1) SIFT2-weighted streamline counts normalised by inverse node volume and squared node radius (connection density per surface area), (2) SIFT2-weighted counts normalised by squared node radius only (preserving size-related variance), and (3) mean streamline length per connection, normalised by squared node radius.

Tractometry provided four additional feature sets. Major 28 white-matter bundles were identified with pyAFQ ^75^, and four tract-level summaries were extracted: mean fractional anisotropy, mean diffusivity, mean streamline length and streamline count. The structural connectivity and tractometry outputs together contributed 16 DWI-derived sets of features.

### Structural MRI: 5 sets of features

We derived five anatomical feature sets from structural MRI outputs processed with FreeSurfer ^80^ version 7.3.2, executed as part of fMRIPrep ^76^. These included cortical thickness, cortical surface area, cortical grey matter volume, subcortical volume, and global brain metrics. We used FreeSurfer’s aparc.stats and aseg.stats files to extract sets of features.

Cortical features were based on the Destrieux atlas ^81^, which partitions the cortex into 148 regions (74 per hemisphere). For each region, we obtained mean thickness, surface area, and grey matter volume.

Subcortical volumes were derived from FreeSurfer’s automatic segmentation using the ASEG atlas ^82^. While the standard ASEG-based set includes 19 commonly used subcortical regions, we retained the full set of 45 labelled regions provided in the aseg.stats output. This includes bilateral subcortical nuclei (e.g., thalamus, putamen, hippocampus), ventricular structures (e.g., lateral, 3rd, 4th, and 5th ventricles), choroid plexus, brainstem, corpus callosum subdivisions, white matter hypointensities, and other midline anatomical structures.

Global brain measures were extracted from the FreeSurfer summary statistics and included total cortical grey matter volume, total white matter volume, subcortical grey matter volume, estimated total intracranial volume (eTIV), and the brain-to-eTIV volume ratio, along with additional summary metrics, resulting in 20 features in this set (see Supplementary Table 38).

### Task-fMRI contrasts (4 sets of features)

We generated four feature sets from task-fMRI contrasts, reflecting changes in BOLD signal associated with events in each task. The BOLD time-series, pre-processed with fMRIPrep ^76^, were analysed on the cortical surface in the symmetric fsLR 91 k space. Detailed descriptions of fMRI preprocessing are provided in Supplementary Methods.

The DLBS includes three fMRI tasks; however, only two, Scenes and Words, were collected consistently across all three waves. We therefore focused on these tasks and excluded the third due to incomplete data. Detailed descriptions can be found in the original publications ^83,84^ or in the “Keys to the Kingdom” dataset documentation^73^. Brief summaries are provided below.

In the Scenes task, which assesses episodic memory, participants viewed landscape images and indicated whether water was present in each scene by pressing a yes/no button. This event-related design presented 96 scenes over three runs, each shown for 3 seconds with variable intertrial intervals. Approximately 20 minutes post-scan, participants completed a recognition task involving 192 images (96 previously seen, 96 novel), classifying each as “high confidence remember,” “low confidence remember,” or “new.” From this task, we computed two orthogonal contrasts: “remembering” (high-confidence imagery minus non-recollected imagery) and “all imagery” (average activation across all presented scenes).

In the Words task, designed to measure semantic processing, participants classified words as living or non-living via button press. This task employed a block design, alternating “easy” (clearly living/non-living) and “hard” (ambiguous) words, such as “speaker,” “virus,” and “sponge.” Participants viewed 128 words, each for 2.5 seconds, interspersed with fixation periods. Two contrasts were derived from this task: a general semantic “task” contrast (combining both difficulty conditions versus fixation) and a “difficulty” contrast (difficult versus easy blocks).

For both the Scenes and Words tasks we applied the same single-subject general linear modelling (GLM) pipeline, implemented in Nilearn ^85^ and Nibabel ^86^ packages. Within each participant every vertex’s time-series was standardized to zero mean and unit variance (z-score), leaving the temporal spectrum otherwise intact.

Nuisance regressors were included following common recommendations to reduce non-neural variance ^87,88^. The design matrix contained 12 head motion parameters, their first derivatives, and a linear trend, five anatomical CompCor ^89^ components extracted from white matter and cerebrospinal fluid to capture physiological noise, and discrete cosine terms for high pass filtering ^87,90^. Event onsets were convolved with the SPM canonical haemodynamic response function and entered alongside the nuisance terms. Models were fit vertex-wise using a first order autoregressive noise model to account for temporal autocorrelation in fMRI time series ^91^. For the Scenes task, beta maps from the three runs were combined using fixed effects with inverse variance weights. The Words task comprised a single run and required no combination.

For each set of task fMRI features, we obtained unthresholded GLM contrasts and parcellated the maps into 379 regions: 360 cortical parcels from the Glasser surface-based atlas ^92^ and 19 subcortical regions from FreeSurfer ASEG ^82^.

### Functional connectivity (FC): 3 sets of features

To complement task contrasts, we generated three additional functional connectivity (FC) feature sets: two from task-based fMRI and one from resting-state fMRI.

Task FC captures background correlations between brain regions during task performance. Unlike GLM-based task contrasts that isolate event-related signal changes, Task FC reflects residual correlations after removing task-locked and confounding effects from the BOLD signal. The preprocessing pipeline followed the same steps used in the task-GLM analysis. The design matrix included HRF-convolved task events and all standardized confounds. For each run, vertex-wise BOLD time series were z-scored and then despiked using a median-based procedure that replaced observations with absolute z-scores above 3.0. Motion outlier volumes were identified from the fMRIPrep confound files as any frame with framewise displacement greater than 0.5 mm or standardised DVARS greater than 1.5, and we additionally censored one preceding and two subsequent volumes.

These censored frames were removed from both the BOLD time series and the design matrix. The nuisance set comprised 12 head motion parameters (three translations and three rotations and their first derivatives), 10 anatomical CompCor components from white matter and cerebrospinal fluid, cosine drift terms and a linear trend. Functional connectivity was computed from the model’s residual parcellated time series. Residuals from this model, representing BOLD signal not explained by task or nuisance regressors, were interpreted as ongoing background activity during the task. For each task (Scenes and Words), we extracted a time series from each of 379 regions, calculated correlations between each region, created a 379 × 379 connectivity matrix and extracted values below the diagonal, yielding two sets of features.

Resting-state FC (rest FC) characterised intrinsic brain connectivity in the absence of task performance. Data processing mirrored the task-based FC pipeline, excluding task regressors. To maintain consistency across participants, only the first resting-state run was analysed due to missing data in approximately one-third of the second runs. This provided a third 379 × 379 connectivity matrix per participant, totalling three FC sets of features.

### Arterial Spin Labeling (ASL): 9 sets of features

We generated nine ASL-derived feature sets from cerebral blood flow (CBF) maps processed using ASLPrep 0.7.5 ^77^. We used the BASIL Bayesian model ^93^ with adaptive spatial regularisation. Sessions with a BASIL quality evaluation index (QEI) below 0.65 were excluded. Further processing and quality-control details are provided in Supplementary Methods.

BASIL produced three tissue-specific CBF maps: grey matter-corrected, white matter-corrected, and an uncorrected global map. Each map was parcellated with the Glasser cortical atlas ^92^ (360 regions); and two compatible subcortical atlases, Tian ^94^ (50 regions) and ASEG ^82^ (19 regions), resulting in nine feature sets. Before modelling, we regressed out three data-quality covariates: BASIL QEI, mean framewise displacement, and ASL-to-T1 registration accuracy using linear regression trained on training data and applied to the test data. Residualised CBF values served as features in all downstream analyses.

### Predictive modelling

We trained predictive, machine learning models to estimate the general cognitive score (g-score) using 37 neuroimaging phenotypes. Each measurement wave was treated as an independent observation, enabling the capture of both within-subject and between-subject variability. Model training employed a nested five-fold cross-validation strategy. To prevent information leakage, all waves from each participant were kept together and randomly assigned to either the training set (80%) or the test set (20%) in each outer fold.

Within each outer training set, a further five-fold split was used for hyperparameter tuning. To select the best hyperparameters, we used the GridSearchCV function of the scikit-learn library. Each inner fold served once as a validation set, while the remaining data formed the corresponding inner training set. Hyperparameters were selected based on validation performance, assessed using the negative mean absolute error. The final model was retrained on the entire outer training set using the selected values before evaluation on the outer test set. Model performance was evaluated using Pearson’s correlation coefficient (r), coefficient of determination (R²), mean absolute error (MAE), and mean squared error (MSE), calculated between predicted and observed g-scores. All phenotype-specific models were trained independently for each feature set.

To examine whether aggregating information across modalities could improve prediction accuracy, we constructed stacked models. After training each phenotype-specific model on the outer training data, we generated predicted scores, which were used as input features for a second-level model. Stacked models were trained exclusively on predictions from the training folds, ensuring the independence of the test data. The same inner cross-validation procedure was used to tune the second-level models.

We defined six stacked models, each corresponding to neuroimaging modalities: (1) task-fMRI, incorporating all task-contrast features; (2) functional connectivity (FC), including both resting-state and task-based connectivity measures; (3) arterial spin labelling (ASL), based on regional cerebral blood flow estimates across different parcellations; (4) diffusion-weighted imaging (DWI), combining structural connectivity and tractometry features; (5) structural MRI (sMRI), aggregating morphometric measures; and (6) a multimodal stacking incorporating features from all phenotypes.

For first-level modelling, we implemented five supervised learning algorithms using Scikit-learn: Elastic Net (EN) ^95^, Partial Least Squares (PLS) ^96^, Random Forest (RF) ^97^, Kernel Ridge Regression (KRR) ^98^, and XGBoost (XGB) ^99^. Stacked models were trained using Random Forest, which accommodates missing values via internal imputation.

In each modelling approach, we conducted exhaustive grid searches over algorithm-specific hyperparameters within a uniform five-fold cross-validation framework. For PLS, we varied the number of latent components from one up to 50 components, selecting the value that minimised mean absolute error. EN models were subjected to a two-dimensional grid search over α and the L1 ratio. The α values spanned 10⁻¹ to 10² (70 logarithmically spaced points) and the L1 ratios ranged from 0 to 1 in 25 equal increments. RF regressors were assessed with 5,000 trees, maximum depths from 1 to 6, and two options for the number of features sampled at each split (square-root and log₂ of the feature count). For XGB, we tuned the learning rate (η at 0.1, 0.2 or 0.3), tree depth (1–6), row subsampling (30□%, 60□% or 100□%), and both L₁ (α) and L₂ (λ) regularisation strengths over zero, 0.5 and 1, again using five-fold CV to balance bias and variance. Finally, in KRR we restricted the kernel to a linear form and searched a wide spectrum of ridge penalties (α from 0 to 20) using mean absolute error as the scoring metric.

### Linear mixed-effects models

We used linear mixed-effects (LME) models on the outer-loop test-set predictions. Each model used the standardised observed g-score (derived from ESEM) as the response variable, denoted g*_observed_*. The explanatory variables included age and/or each MRI marker, or the predicted g-score, derived from 37 neuroimaging phenotypes, as well as six modality-specific and multimodal stacking models generated using the best-performing algorithm.

Age was grand-mean centred, and two components were created to separate between-person and within-person variability: age*_mean_* (an individual’s average age across waves), representing between-person effects, and age*_dev_*(the deviation from age*_mean_* at each wave), representing within-person effects ^100^.

Each MRI marker was standardised across all observations. Similar to age, two components were computed: *g_pred_mean_* (each person’s average predicted g-score across waves), representing between-person effects, and *g_pred_dev_* (the person-centred deviation from *g_pred_mean_*), representing within-person effects ^100^.

### The relationship between g-score and age

The first two LME models estimated the relationship between g-score and age. The first model, the age-additive model, included age*_mean_* and age*_dev_* as fixed effects and had a random intercept, using the following lme4 syntax:

### Age-additive model

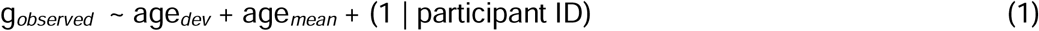

The second model, the age-interactive model, extended the age-additive model by including an interaction term between age*_mean_* and age*_dev_*. The interactive model tested whether the within-person effect of age on cognitive functioning (age*_dev_*) varied depending on a person’s average age (age*_mean_*). Given the wide age range in the sample, it is plausible that age-related cognitive decline within individuals is steeper for older adults than for younger adults. Importantly, adding the interaction term makes the main effects of age*_mean_* and age*_dev_* conditional. For instance, the main effect of age*_dev_* now reflects the effect of within-person age change when age*_mean_* is zero, that is, at the sample’s average age. We used the following lme4 syntax:

### Age-interactive model

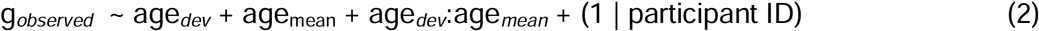

We compared these two age models using the likelihood ratio test. If having the interactive term improved model fit, we would keep the age-interactive model.

### Variance decomposition

The third LME model, the neuroimaging-only model, allowed us to test the extent to which MRI markers explained within-person versus between-person variance in the observed g-score. This model included both between-person (*g_pred_mean_*) and within-person (*g_pred_dev_*) MRI markers as fixed effects and had random intercepts, using the following lme4 syntax

### Neuroimaging-only model

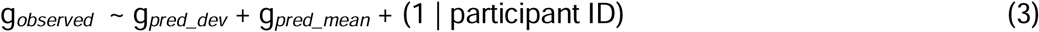

To assess how much within-person and between-person variance in cognitive functioning was explained by MRI markers, we applied variance decomposition to the neuroimaging-only models ^38^. The observed g-scores were partitioned into within-person (level-1) and between-person (level-2) variance components:

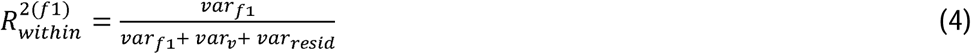

Where var_f1_ represents variance explained by the first-level fixed effect (i.e., the within-person MRI marker), var_v_ captures variance associated with random-slope effects (equals zero in our model), and var_resid_ is the level-1 residual variance.

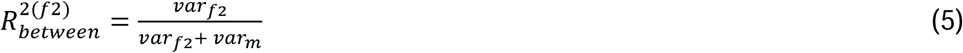

Where var_f2_ represents the variance explained by the second-level fixed effect (i.e., the between-person MRI marker), and var_m_ is the random intercept variance.

These coefficients provide separate within-and between-person estimates of explanatory power for each MRI-based marker.

### Commonality analyses

The fourth model, the age-plus-neuroimaging model, included both age variables and each MRI marker as explanatory variables, allowing simultaneous estimation of age-related and neuroimaging-related effects and quantification of their unique and shared contributions to cognitive functioning.

Because the age interaction term improved model fit in the age-interactive model relative to the age-additive model (see Results), we included age*_dev_*, age*_mean_*, and their interaction (age*_dev_*:age*_mean_*) as fixed effects for age, along with *g_pred_dev_* and *g_pred_mean_* as fixed effects for the MRI markers. The lme4 syntax was:

### Age-plus-neuroimaging model

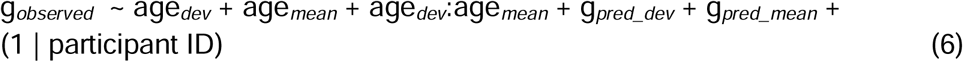

To quantify the extent to which MRI markers explained age-related variation in cognitive functioning, we performed a commonality analysis^41,42^. This method examines changes in marginal R², the proportion of variance explained by the fixed effects, when each fixed-effect explanatory variable is included or removed.

Comparing these changes allows partitioning the variance in g*_observed_* into: a) unique variance, attributable solely to age*_dev_*, age*_mean_*, age*_dev_*:age*_mean_*, g*_pred_dev_*, or g*_pred_mean_*, and b) shared variance, attributable to combinations of these predictors.

Using the commonality analysis, we derived four proportions:

a) Within-person age-related variance explained by within-person MRI markers: Shared variance between age*_dev_*, and g*_pred_dev_* across all combinations of fixed-effect variables, divided by the total variance of age*_dev_*.
b) Between-person age-related variance explained by between-person MRI markers: Shared variance between age*_mean_* and g*_pred_mean_* across all combinations of fixed-effect variables, divided by the total variance of age*_mean_*.
c) Ag e-interaction variance explained by within-person MRI markers: Shared variance between age*_dev_*:age*_mean_* and g*_pred_dev_* across all combinations of fixed-effect variables, divided by the total variance of age*_dev_*:age*_mean_*.
d) Age-interaction variance explained by between-person MRI markers: Shared variance between age*_dev_*:age*_mean_* and g*_pred_mean_* across all combinations of fixed-effect variables, divided by the total variance of age*_dev_*:age*_mean_*.

All mixed-effects analyses were conducted in R (version 4.4.1), using tidyverse (v2.0.0) for data preparation, lme4 (v1.1-37) for model fitting, r2mlm (v0.3.8) for variance decomposition, and glmm.hp (v0.1.8) for commonality analysis.

## Data availability

We used publicly available DLBS 1.2.0 data (doi:10.18112/openneuro.ds004856.v1.2.0) provided by the Dallas Lifespan Brain Study (https://openneuro.org/datasets/ds004856/versions/1.2.0).

## Code availability

The codes for analyzing the data are available at https://github.com/HAM-lab-Otago-University/Multimodal-MRI-prediction-Dallas-dataset

