## Supplementary methods for "Multimodal MRI prediction of cognitive functioning across the lifespan: separating between-person differences from within-person changes"

### DWI

*Preprocessing*

Preprocessing was performed using QSIPrep 1.0.1.dev0+gee9aa2e.d20250115 (Cieslak et al. 2021), which is based on Nipype 1.9.1 [Gorgolewski et al. (2011); Gorgolewski et al. (2018); RRID:SCR_002502].

The anatomical reference image was reoriented into AC-PC alignment via a 6-DOF transform extracted from a full Affine registration to the MNI152NLin2009cAsym template. A full nonlinear registration to the template from AC-PC space was estimated via symmetric nonlinear registration (SyN) using antsRegistration. Brain extraction was performed on the T1w image using SynthStrip (Hoopes et al. 2022) and automated segmentation was performed using SynthSeg (Billot, Greve, et al. 2023, @synthseg2) from FreeSurfer version 7.3.1.

DWI data were denoised using the Marchenko-Pastur PCA method implemented in dwidenoise (Tournier et al. 2019; Veraart et al. 2016; Veraart, Fieremans, and Novikov 2016; Cordero-Grande et al. 2019) with an automatically-determined window size of 3 voxels. After MP-PCA, Gibbs ringing was removed using MRtrix3 (Tournier et al. 2019; Kellner et al. 2016). Any images with a b-value less than 100 s/mm^2 were treated as a b=0 image. The mean intensity of the DWI series was adjusted so all the mean intensity of the b=0 images matched across each separate DWI scanning sequence. B1 field inhomogeneity was corrected using dwibiascorrect from MRtrix3 with the N4 algorithm (Tustison et al. 2010) after corrected images were resampled.

FSL (version None)’s eddy was used for head motion correction and Eddy current correction (Andersson and Sotiropoulos 2016). Eddy was configured with a q-space smoothing factor of 10, a total of 5 iterations, and 1000 voxels used to estimate hyperparameters. A quadratic first level model and a linear second level model were used to characterize Eddy current-related spatial distortion. q-space coordinates were forcefully assigned to shells. Field offset was attempted to be separated from subject movement. Shells were aligned post-eddy. Eddy’s outlier replacement was run (Andersson et al. 2016). Data were grouped by slice, only including values from slices determined to contain at least 250 intracerebral voxels. Groups deviating by more than 4 standard deviations from the prediction had their data replaced with imputed values. Final interpolation was performed using the jac method.

Several confounding time-series were calculated based on the preprocessed DWI: framewise displacement (FD) using the implementation in Nipype (following the definitions by Power et al. 2014). The head-motion estimates calculated in the correction step were also placed within the corresponding confounds file. Slicewise cross correlation was also calculated. The DWI time-series were resampled to ACPC, generating a preprocessed DWI run in ACPC space with 1.8mm isotropic voxels.

*Tractography*

Diffusion-weighted images were preprocessed and reconstructed using QSIRecon (version 1.0.1.dev0+gf78c888; Cieslak et al., 2021), which operates within the Nipype neuroimaging framework (v1.9.1; Gorgolewski et al., 2011, 2018). Given the single-shell nature of our diffusion acquisition, we employed single-shell three-tissue constrained spherical deconvolution (SS3T-CSD) to estimate fibre orientation distributions (FODs). This approach simultaneously models white matter (WM), grey matter (GM), and cerebrospinal fluid (CSF) compartments, enabling the resolution of crossing fibres from single-shell data (Dhollander & Connelly, 2016; Dhollander et al., 2016). Multi-tissue response functions for each tissue type were estimated using the Dhollander algorithm implemented in MRtrix3.

To generate biologically plausible tractograms, we applied anatomically constrained tractography (ACT). Each participant’s T1-weighted anatomical image was segmented into tissue classes using a hybrid surface-volume segmentation (hsvs) pipeline, which informed streamline propagation and termination criteria. This ensured that streamlines were anatomically consistent, avoiding entry into non-white matter regions (Smith et al., 2012). Probabilistic streamline tracking was performed using the iFOD2 algorithm in MRtrix3, with seeding throughout the white matter FODs. Streamlines were terminated upon reaching a minimum FOD amplitude threshold or upon leaving the white matter compartment. Length constraints and ACT-based filters were used to exclude implausibly short, long, or looping trajectories.

*Structural Connectivity*

Whole-brain tractography results were used to construct atlas-based structural connectivity matrices. For each participant the brain was parcellated with four atlases: three whole-brain Schaefer-based 4S variants (4S156, 4S256 and 4S456) and the 333-parcel cortical Gordon atlas. For each atlas, parcels defined network nodes, and edges were quantified as the number of streamlines connecting each pair. This yielded a set of connectivity matrices per participant summarising pairwise structural connectivity.

From each atlas, we extracted three automatically generated connectivity matrices for use in downstream machine learning analyses. The first (sift_invnodevol_radius2_countconnectome) applied SIFT2-optimised streamline weights, scaled by the inverse mean node volume and the squared node radius. This adjustment reduces bias toward larger parcels and normalises by cortical surface area, providing an estimate of connection density per unit surface. The second matrix (sift_radius2_countconnectome) also used SIFT2 weights and scaled by squared node radius but omitted the volume correction, thereby preserving size-related variance in connectivity strength. The third matrix (radius2_meanlengthconnectome) used mean streamline length per connection, scaled by squared node radius. Length weighting corrects for tractography’s bias toward shorter streamlines and provides an estimate of conduction distance between regions. Together, these three matrix types across four atlases produced twelve feature sets used in predictive modelling.

*Tractometry*

In addition, we incorporated the Python Automated Fiber Quantification (pyAFQ) to identify major white matter tracts from whole-brain tractography. PyAFQ identifies major 28 white matter bundles based on predefined anatomical criteria using a combination of waypoint ROIs and orientation filters. Only streamlines meeting bundle-specific inclusion criteria were retained, others were excluded as non-bundle or artifactual.

The full list of white matter bundles included: Callosum Anterior Frontal, Callosum Motor, Callosum Occipital, Callosum Orbital, Callosum Posterior Parietal, Callosum Superior Frontal, Callosum Superior Parietal, Callosum Temporal, Left Anterior Thalamic, Left Arcuate, Left Cingulum Cingulate, Left Corticospinal, Left Inferior Fronto-occipital, Left Inferior Longitudinal, Left Posterior Arcuate, Left Superior Longitudinal, Left Uncinate, Left Vertical Occipital, Right Anterior Thalamic, Right Arcuate, Right Cingulum Cingulate, Right Corticospinal, Right Inferior Fronto-occipital, Right Inferior Longitudinal, Right Posterior Arcuate, Right Superior Longitudinal, Right Uncinate, Right Vertical Occipital.

Once bundles were identified, tract profiles of diffusion properties were extracted along each tract’s trajectory. For each bundle, fractional anisotropy (FA) and mean diffusivity (MD) values were sampled at equidistant nodes along the tract (from one terminus to the other) to represent a tract-specific profile (series of measurements). We extracted tract-level summary metrics including the mean FA and MD, the total number of streamlines per tract, and the mean streamline length for each bundle. These summary metrics served as four additional sets of features covering DWI modality.

### Structural MRI

*Preprocessing*

The T1-weighted (T1w) images were corrected for intensity non-uniformity (INU) with N4BiasFieldCorrection (Tustison et al. 2010), distributed with ANTs 2.5.3 (Avants et al. 2008, RRID:SCR_004757), and used as T1w-reference throughout the workflow. The T1w-reference was then skull-stripped with a Nipype implementation of the antsBrainExtraction.sh workflow (from ANTs), using OASIS30ANTs as target template. Brain tissue segmentation of cerebrospinal fluid (CSF), white-matter (WM) and gray-matter (GM) was performed on the brain-extracted T1w using fast (FSL (version unknown), RRID:SCR_002823, Zhang, Brady, and Smith 2001). Brain surfaces were reconstructed using recon-all (FreeSurfer 7.3.2, RRID:SCR_001847, Dale, Fischl, and Sereno 1999), and the brain mask estimated previously was refined with a custom variation of the method to reconcile ANTs-derived and FreeSurfer-derived segmentations of the cortical gray-matter of Mindboggle (RRID:SCR_002438, Klein et al. 2017). A T2-weighted image was used to improve pial surface refinement. Brain surfaces were reconstructed using recon-all (FreeSurfer 7.3.2, RRID:SCR_001847, Dale, Fischl, and Sereno 1999), and the brain mask estimated previously was refined with a custom variation of the method to reconcile ANTs-derived and FreeSurfer-derived segmentations of the cortical gray-matter of Mindboggle (RRID:SCR_002438, Klein et al. 2017). Volume-based spatial normalization to two standard spaces (MNI152NLin2009cAsym, MNI152NLin6Asym) was performed through nonlinear registration with antsRegistration (ANTs 2.5.3), using brain-extracted versions of both T1w reference and the T1w template. The following templates were were selected for spatial normalization and accessed with TemplateFlow (24.2.0, Ciric et al. 2022): ICBM 152 Nonlinear Asymmetrical template version 2009c [Fonov et al. (2009), RRID:SCR_008796; TemplateFlow ID: MNI152NLin2009cAsym], FSL’s MNI ICBM 152 non-linear 6th Generation Asymmetric Average Brain Stereotaxic Registration Model [Evans et al. (2012), RRID:SCR_002823; TemplateFlow ID: MNI152NLin6Asym]. Grayordinate “dscalar” files containing 91k samples were resampled onto fsLR using the Connectome Workbench (Glasser et al. 2013).

### Functional MRI

*Preprocessing*

Results included in this manuscript come from preprocessing performed using fMRIPrep 24.1.1 (Esteban et al. (2019); Esteban et al. (2018); RRID:SCR_016216), which is based on Nipype 1.8.6 (K. Gorgolewski et al. (2011); K. J. Gorgolewski et al. (2018); RRID:SCR_002502).

For each of the BOLD runs found per subject (across all tasks and sessions), the following preprocessing was performed. First, a reference volume was generated, using a custom methodology of fMRIPrep, for use in head motion correction. Head-motion parameters with respect to the BOLD reference (transformation matrices, and six corresponding rotation and translation parameters) are estimated before any spatiotemporal filtering using mcflirt (FSL , Jenkinson et al. 2002). The BOLD reference was then co-registered to the T1w reference using bbregister (FreeSurfer) which implements boundary-based registration (Greve and Fischl 2009). Co-registration was configured with six degrees of freedom. The aligned T2w image was used for initial co-registration.Several confounding time-series were calculated based on the preprocessed BOLD: framewise displacement (FD), DVARS and three region-wise global signals. FD was computed using two formulations following Power (absolute sum of relative motions, Power et al. (2014)) and Jenkinson (relative root mean square displacement between affines, Jenkinson et al. (2002)). FD and DVARS are calculated for each functional run, both using their implementations in Nipype (following the definitions by Power et al. 2014). The three global signals are extracted within the CSF, the WM, and the whole-brain masks. Additionally, a set of physiological regressors were extracted to allow for component-based noise correction (CompCor, Behzadi et al. 2007). Principal components are estimated after high-pass filtering the preprocessed BOLD time-series (using a discrete cosine filter with 128s cut-off) for the two CompCor variants: temporal (tCompCor) and anatomical (aCompCor). tCompCor components are then calculated from the top 2% variable voxels within the brain mask. For aCompCor, three probabilistic masks (CSF, WM and combined CSF+WM) are generated in anatomical space. The implementation differs from that of Behzadi et al. in that instead of eroding the masks by 2 pixels on BOLD space, a mask of pixels that likely contain a volume fraction of GM is subtracted from the aCompCor masks. This mask is obtained by dilating a GM mask extracted from the FreeSurfer’s aseg segmentation, and it ensures components are not extracted from voxels containing a minimal fraction of GM. Finally, these masks are resampled into BOLD space and binarized by thresholding at 0.99 (as in the original implementation). Components are also calculated separately within the WM and CSF masks. For each CompCor decomposition, the k components with the largest singular values are retained, such that the retained components’ time series are sufficient to explain 50 percent of variance across the nuisance mask (CSF, WM, combined, or temporal). The remaining components are dropped from consideration. The head-motion estimates calculated in the correction step were also placed within the corresponding confounds file. The confound time series derived from head motion estimates and global signals were expanded with the inclusion of temporal derivatives and quadratic terms for each (Satterthwaite et al. 2013). Frames that exceeded a threshold of 0.5 mm FD or 1.5 standardized DVARS were annotated as motion outliers. Additional nuisance timeseries are calculated by means of principal components analysis of the signal found within a thin band (crown) of voxels around the edge of the brain, as proposed by (Patriat, Reynolds, and Birn 2017). The BOLD time-series were resampled onto the left/right-symmetric template “fsLR” using the Connectome Workbench (Glasser et al. 2013). Grayordinates files (Glasser et al. 2013) containing 91k samples were also generated with surface data transformed directly to fsLR space and subcortical data transformed to 2 mm resolution MNI152NLin6Asym space. All resamplings can be performed with a single interpolation step by composing all the pertinent transformations (i.e. head-motion transform matrices, susceptibility distortion correction when available, and co-registrations to anatomical and output spaces). Gridded (volumetric) resamplings were performed using nitransforms, configured with cubic B-spline interpolation.

### ASL

*Preprocessing*

Arterial spin-labeled MRI images were preprocessed using ASLPrep 0.7.5 (Adebimpe et al. 2022, 2023), which is based on fMRIPrep (Esteban et al. (2019); Esteban et al. (2020); RRID:SCR_016216) and Nipype 1.8.6 (Gorgolewski et al. 2011).

For each subject, a single ASL run was identified across all tasks and sessions and subjected to the following preprocessing. A reference volume was first generated using ASLPrep’s custom method, to support subsequent head motion correction. Head motion parameters were then estimated using FSL’s mcflirt tool (Jenkinson et al., 2002). To avoid confounding intensity differences between contrasts with motion artefacts, motion correction was performed separately for each volume type (Wang et al., 2008). The motion parameters from each volume type were then concatenated, and ASLPrep re-computed the relative root mean squared deviation.

ASLPrep calculated cerebral blood flow (CBF) from the single-delay PCASL using a single-compartment general kinetic model (Buxton et al. 1998). As no calibration images or provided M0 estimate was available for the ASL scan, the control volumes used as a substitute. The control volumes in the ASL scans were smoothed with a Gaussian kernel (FWHM=5.0 mm) and the average control image was calculated and scaled by 1.

Structural Correlation based Outlier Rejection (SCORE) algorithm was applied to the CBF timeseries to discard CBF volumes with outlying values (Dolui et al. 2017) before computing the mean CBF. Following SCORE, the Structural Correlation with RobUst Bayesian (SCRUB) algorithm was applied to the CBF maps using structural tissue probability maps to reweight the mean CBF (Dolui et al. 2017; Dolui, Wolk, et al. 2016).

CBF was also computed with Bayesian Inference for Arterial Spin Labeling (BASIL) (Michael A. Chappell et al. 2009), as implemented in FSL None. BASIL computes CBF using a spatial regularization of the estimated perfusion image and additionally calculates a partial-volume corrected CBF image (M. A. Chappell et al. 2011). Several confounding timeseries were calculated, including both framewise displacement (FD) and DVARS. FD and DVARS are calculated using the implementations in in Nipype (following the definition by (Power et al. 2014)) for each ASL run. ASLPrep summarizes in-scanner motion as the mean framewise displacement and relative root-mean square displacement.

Parcellated CBF estimates were extracted for the following atlases: the Schaefer Supplemented with Subcortical Structures (4S) atlas (Schaefer et al. 2018; Pauli, Nili, and Tyszka 2018; King et al. 2019; Najdenovska et al. 2018; Glasser et al. 2013) at 10 different resolutions (156, 256, 356, 456, 556, 656, 756, 856, 956, and 1056 parcels), the Glasser atlas (Glasser et al. 2016), the Gordon atlas (Gordon et al. 2016), the Tian subcortical atlas (Tian et al. 2020), and the HCP CIFTI subcortical atlas (Glasser et al. 2013). In cases of partial coverage, either uncovered voxels (values of all zeros or NaNs) were ignored (when the parcel had >50.0% coverage) or the whole parcel was set to zero (when the parcel had <50.0% coverage).

The quality evaluation index (QEI) was computed for each CBF map (Dolui, Wolf, et al. 2016). QEI is based on the similarity between the CBF and the structural images, the spatial variability of the CBF image, and the percentage of grey matter voxels containing negative CBF values. All resampling in ASLPrep uses a single interpolation step that concatenates all transformations. Gridded (volumetric) resampling was performed using antsApplyTransforms, configured with Lanczos interpolation to minimize the smoothing effects of other kernels (Lanczos 1964). Many internal operations of ASLPrep use Nilearn 0.11.1 (Abraham et al. 2014), NumPy (Harris et al. 2020), and SciPy (Virtanen et al. 2020).

*Features selection*

The default ASLPrep workflow derives a perfusion-weighted timeseries by subtracting label images from control images across repeated acquisitions. These difference volumes are averaged into a mean cerebral blood flow (CBF) map and evaluated using the quality evaluation index (QEI), which ranges from 0 to 1. QEI quantifies image quality based on anatomical similarity, spatial smoothness, and the proportion of negative voxels in grey matter. In our dataset, only 2 % of sessions reached a QEI of at least 0.65, indicating that standard mean CBF maps, computed without spatial regularisation, were typically of low quality.

To improve the reliability of CBF estimates, we enabled BASIL (Bayesian Inference for Arterial Spin Labelling) within ASLPrep. BASIL fits a generative Bayesian model that incorporates adaptive spatial regularisation, which assumes that neighbouring voxels within brain tissue have similar CBF values (Chappell et al., 2009, 2011). This regularisation suppresses voxel-wise noise and generates a single spatially smoothed, temporally averaged CBF map per session. ASLPrep applies the same QEI framework to this map, reporting the score as qei_cbf_basil. In our sample, 76 % of sessions exceeded the 0.65 QEI threshold when evaluated on BASIL-derived maps, suggesting that spatial regularisation substantially enhanced image quality. Sessions falling below this threshold were excluded from further analysis.

Although no universal QEI cutoff has been adopted in the literature, previous studies have used thresholds ranging from 0.4 to 0.9 depending on dataset quality and analysis goals (Ketcherside et al., 2020; Yu et al., 2025; Rashid et al., 2023). We adopted a slightly relaxed version of the 0.70 threshold used by Ketcherside et al., setting a cutoff at 0.65 to retain average-quality scans while excluding those of clearly insufficient quality.

BASIL, as implemented in ASLPrep, produced tissue-specific CBF estimates for grey matter (GM), white matter (WM), and the whole brain by applying partial volume correction. This correction accounts for the partial volume effects that arise when individual voxels contain a mixture of tissue types, improving the accuracy of perfusion quantification within each tissue class. ASLPrep outputs three corresponding CBF maps: one corrected for GM, one for WM, and one uncorrected (reflecting global perfusion without tissue-type specificity). Each of these three CBF maps was parcellated to extract regional mean CBF values. Due to incomplete coverage or missing data in some atlases, we restricted our analysis to three parcellations that provided consistent coverage across sessions: the Glasser cortical atlas, the Tian subcortical atlas, and the HCP subcortical parcellation.These left us with nine feature sets, corresponding to the combination of three CBF map types and three parcellation atlases.

Prior to predictive modeling, we applied linear regression to remove the influence of several data-quality covariates on the regional CBF measures. In particular, we regressed out BASIL QEI, mean framewise displacement, and the ASL-to-T1 coregistration dice coefficient. All confound variables were z-scored, the regression model was fitted on the training set and applied to the test set, and the resulting residualised CBF values were used as features to predict participants’ cognitive G-scores in subsequent analyses.
